# ZipSeq : Barcoding for Real-time Mapping of Single Cell Transcriptomes

**DOI:** 10.1101/2020.02.04.932988

**Authors:** Kenneth H. Hu, John P. Eichorst, Chris S. McGinnis, David M. Patterson, Eric D. Chow, Kelly Kersten, Stephen C. Jameson, Zev J. Gartner, Arjun A. Rao, Matthew F. Krummel

**Author notes:** Corresponding Author: Matthew F. Krummel, Ph.D., 513 Parnassus Avenue, HSW 512, San Francisco, CA 94143-0511.

## Abstract

Spatial transcriptomics seeks to integrate single-cell transcriptomic data within the 3-dimensional space of multicellular biology. Current methods use glass substrates pre-seeded with matrices of barcodes or fluorescence hybridization of a limited number of probes. We developed an alternative approach, called ‘ZipSeq’, that uses patterned illumination and photocaged oligonucleotides to serially print barcodes (Zipcodes) onto live cells within intact tissues, in real-time and with on-the-fly selection of patterns. Using ZipSeq, we mapped gene expression in three settings: in-vitro wound healing, live lymph node sections and in a live tumor microenvironment (TME). In all cases, we discovered new gene expression patterns associated with histological structures. In the TME, this demonstrated a trajectory of myeloid and T cell differentiation, from periphery inward. A variation of ZipSeq efficiently scales to the level of single cells, providing a pathway for complete mapping of live tissues, subsequent to real-time imaging or perturbation.

## INTRODUCTION

Single cell RNA Sequencing (scRNA-Seq) combined with other multimodal analyses such as surface epitope labeling and repertoire analysis have revealed previously unappreciated heterogeneity within cell populations. This approach has been especially useful in immunology given the diversity of immune cell types and the microenvironments they experience.^1^. Yet scRNA-Seq studies lose information on the spatial context where a given single cell transcriptome was localized^2^. Conventional microscopy localizes cells and molecules in space, however, it is limited in the parameters it can detect. Even with high dimensional imaging techniques (MIBI, CODEX), probes must be selected a priori^3, 4^. In order to better understand specifically how cellular transcriptional heterogeneity is influenced by the local environment and vice versa in a discovery-based, unbiased approach, it becomes necessary to link high-dimensional scRNA-Seq data to the spatial dimensions and real-time phenotypical analyses that microscopy affords.

To couple conventional microscopy of live tissues with transcriptomics, we needed a means to demarcate multiple manually defined regions-of-interest (ROI’s) in real-time, in both mouse and human tissues. While excellent for some applications, we decided against grid-based approaches^5–7^, which average together gene expression within a region and are defined prior to imaging on fixed tissue sections. Instead, we sought to use existing scRNA-Seq workflows and to develop a method for ‘printing’ a DNA barcode onto live cells in a spatially defined manner, which can be read-out during sequencing^8^. We accomplished this by initially coating a base DNA oligo onto cells in a tissue and –through photocaging—we could control hybridization of subsequent DNA strands in a light- and thus spatially-restricted manner^9, 10^. The resultant barcodes then provided a connection to those user defined regions.

## RESULTS

We generated a photo-uncaging system that allowed light-based printing of DNA barcodes onto the surface of cells. A double-stranded piece of DNA was attached to cells, either by a high affinity antibody (e.g. anti-CD45, for pan-immune cells) or via stable lipid insertion (e.g. lignoceric acid) into the membrane^11^. This double stranded “anchor strand” contained a 16 bp overhang sequence (termed “O1”) that is blocked at four sites along its length using 6-nitropiperonyloxylmethyl (NPOM) conjugated to thymidine, and thus is unable to participate in base-pairing^9^. However, following local illumination with 365 nm light to release the cages, a readout oligonucleotide strand, termed a “Zipcode” or ‘ZC’ strand can hybridize to O1 (**Fig. 1A**). This annealed Zipcode terminates in a polyA sequence and an Illumina Read 2 Sequence which then allows for poly-dT based amplification during library construction^8^. We first demonstrated light-based control of hybridization for two readout strands, by marking two separated populations of primary mouse CD4 and CD8 T cells and demonstrating excellent concordance of populations and the associated Zipcodes (**Fig. S1**).

**Figure 1:**
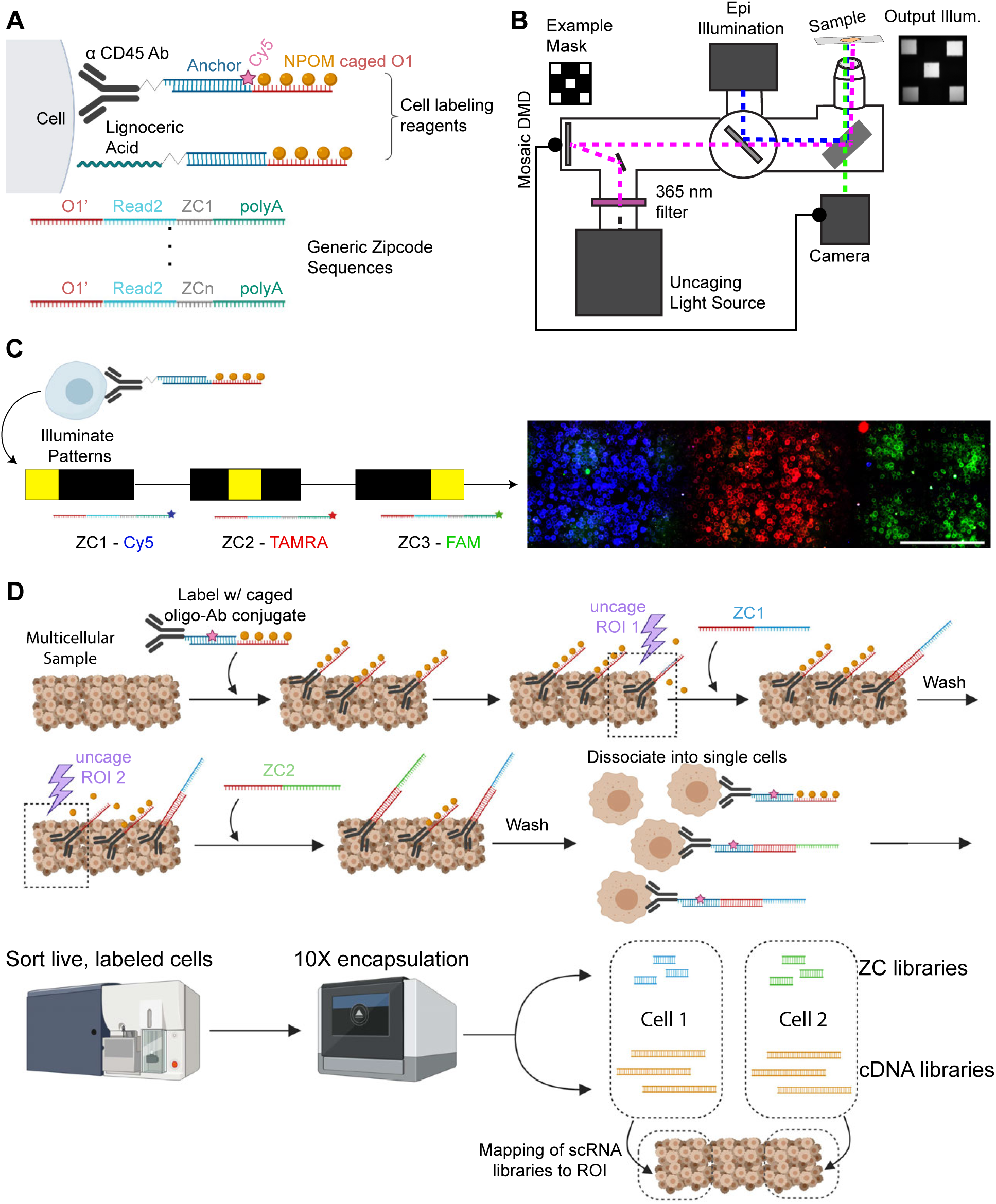
**(a)** Schematic of oligonucleotide sequences and labeling moieties used in this paper. Both lipid and antibody are covalently conjugated to an ‘anchor’ sequence. Meanwhile a caged strand consisting of 4 photocaging groups on an overhang sequence linked to the reverse complement of the anchor strand can hybridize with the Ab or lipid DNA conjugate prior to labeling cells. Readout strand or ‘Zipcodes’ consist of a reverse complement sequence to the caged overhang sequence 1 or 2, followed by a partial Illumina Small RNA Read 2 sequence for downstream amplification. In addition, each Zipcode strand bears a 8bp barcode and a 28 polyA sequence for capture by poly dT primers during reverse transcription. **(b)** A microscope light path for simultaneous imaging and photo-uncaging of a sample. Spatially directed photo-uncaging is accomplished through directing light from a mercury arc-lamp onto a Mosaic DMD with an 800 × 600 micromirror array in plane with the sample. The sample can be simultaneously imaged using epi-fluorescent excitation. In the imaging software, a user defined ROI is converted into a mask which is reflected in the micromirror array. This spatially patterned light is then directed through the microscope and objective onto the sample. An example mask is shown with the resulting illumination pattern visualized on a mirrored slide. **(c)** Illustration of proof-of-concept demonstrating ability to spatially control hybridization of fluorescently labelled oligonucleotides. Briefly, a monolayer of primary mouse CD8 T cells was plated, labeled w/ the anti-CD45 Ab-DNA conjugate with the caged overhang O1. The leftmost square ROI of 300 µm size was illuminated with 365 nm light and the first Zipcode was added and allowed to hybridize. Following wash steps, the process was repeated 2x at other positions with distinct Zipcode-fluorophore combinations resulting in 3 defined regions. Scale bar = 200 µm **(d)** Schematic for workflow for labeling two regions of interest in a tissue section beginning with labeling of cells in a dish/in tissue with appropriate labeling moiety hybridized to a strand bearing the photocaged ssDNA overhang. Manual delineation of an ROI is followed by a pulse of UV illumination. Addition of a readout strand or Zipcode 1 allows for labeling of uncaged overhangs i.e. cells within ROI. Following washout of this Zipcode 1, the process is repeated for ROI #2. Cells are then harvested or dissociated from tissue, (FACS-)sorted for labelled cells, and passed to the 10X Chromium Controller for encapsulation and reverse transcription.

We used a digital micromirror device (DMD) to control the spatial pattern of 365 nm light in a plane conjugate with the image plane of a conventional Zeiss widefield microscope (**Fig. 1B**). The nature of the DMD allows individual pixels to be illuminated down to approximately the resolution limit of the objective used.

To demonstrate specific printing of a collection of Zipcodes, we plated CD8 mouse T cells, labeled with caged anti-CD45 Ab-DNA anchor strands in a chamber slide and used Zipcode strands that were labeled with one of three fluorophores to visually track spatially-controlled annealing. Following three rounds of: (1. Patterned illumination 2. Zipcode addition 3. Washing), we obtained clear delineation of 3 regions, showing the linear scaling of resolution and number of rounds (**Fig. 1C**). Individual cells that floated through the imaging field are likely the cause of deviations from the expected labeling scheme. Taking these two proofs of concept together, a schematic of an idealized workflow is depicted in **Fig. 1D**. Note that an internal fluorophore such as Cy5 can be incorporated into the anchor sequence or a terminal Zipcode, allowing for enrichment of labeled cells before encapsulation.

### Defining Spatially-Segregated Motility and Cell Division Programs in Wound Healing

We applied ZipSeq to study spatially-defined transcriptional programs in a well defined model of wound healing in a monolayer of NIH/3T3 fibroblasts. 12 hours after ‘wounding’, we performed live-imaging of the wound edge (**Fig. 2A**). We then used the lignoceric acid conjugated photocaged anchor strand (**Fig. 1A**) to first label all cells and then illuminated a band 0-200 µm from the wound edge (“Front”) and added Zipcode 1. Following a brief incubation, we washed this oligonucleotide out, and illuminated another band 200-400 µm away from the wound edge (“Rear”) followed by Zipcode 2 addition (**Fig. 2B**). We then dissociated the monolayer into single cells and subjected the cells to a 10X scRNA-Seq pipeline.

**Figure 2:**
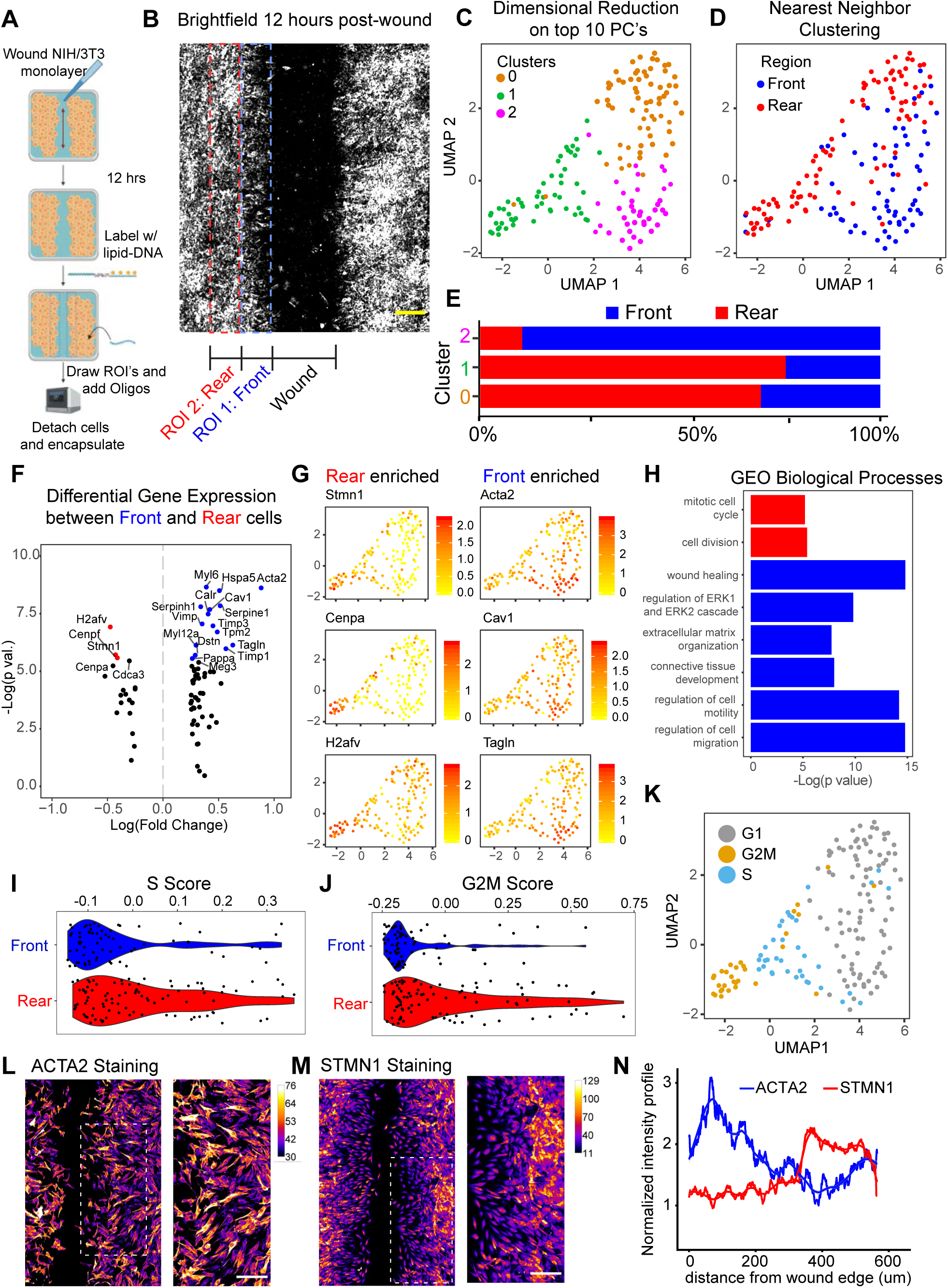
ZipSeq Mapping of a Live Cell Monolayer Following Wounding. **(a)** Experimental setup. NIH/3T3 cells were plated 48 hours prior to imaging and allowed to reach confluency. 12 hours prior to imaging, a pipette tip was used to cleanly scrape away a band. The wound was imaged after 12 hours and ROI’s defined. Cells were then labeled with a lipid-oligo conjugate and then uncaged in a series of vertical bands alternating with Zipcode addition. Harvested cells were then passed into the modified 10X workflow. **(b)** Brightfield of wound 12 hours post-wounding with ROI’s overlaid. Two vertical bands of 200 µm width were drawn with increasing distance from the wound edge (0-200 µm) and (200-400 µm), referred to as ‘front’ and ‘rear’ respectively for illumination and zipcoding. Scale bar = 200 µm. **(c)** UMAP representation of labeled cells with majority ZC identity overlaid. UMAP was calculated using the top 10 principal components. **(d)** UMAP representation of labeled cells with cluster overlay. Clusters calculated using Seurat’s built-in SNN based clustering algorithm. **(e)** Percentage of cells belonging to either Front or Rear populations within each cluster as defined in **d**. **(f)** Volcano plot from differential expression analysis between Front and Rear cells. Colored points represent genes with an adjusted p-value (Bonferroni corrected) <0.05. **(g)** Feature plots overlaid on UMAP representation for 3 selected genes from DE analysis enriched in either Front (*Acta2*, *Cav1*, and *Tagln*) or Rear cells (*Stmn1*, *Cenpa*, *H2afv*). Color scale indicates Log-Normalized gene read counts. **(h)** Hits from DE analysis were passed through GO analysis. Significantly enriched biological processes shown with -Log(p-value) (Bonferroni corrected). Violin plots for **(i)** S phase and **(j)** G2M phase signature score for Front and Rear cells. Gene lists used are shown in Methods. **(k)** Assignment to cell cycle phase (S, G2M or G1 phase) based on the signature scores calculated in (i) and (j). Immunofluorescence imaging of fixed NIH/3T3 cells 12 hours post-wounding stained for either **(l)** ACTA2 or **(m)** STMN1. Fire LUT from ImageJ applied. Zoomed in insets shown for indicated regions. Scale bar = 100 µm. **(N)** Line plot with quantification of mean fluorescence intensity vs. distance from edge. IF images from (l and m) were first masked for pixels belonging to cells vs. background. Then in-cell pixels within vertical bands stepping away from the wound were averaged to create the indicated line-scan profiles with a smoothed fit applied.

During analysis, we removed the cells with low Zipcode counts (i.e. from neither illuminated zone) from the analysis and used the ratio of Zipcode 1 to 2 counts to determine whether a cell derived from the Front region versus the Rear. We then performed unsupervised, nearest neighbor clustering combined with UMAP dimensional reduction to visualize labeled cells in transcriptional space and identified three clusters (**Fig. 2C**). When we overlaid the ZipSeq region calls, a clear partitioning between Front and Rear cells was observed in transcriptional space (**Fig. 2D**), with cluster 2 highly enriched for Front cells and clusters 0 and 1 relatively enriched for Rear cells (**Fig. 2E**).

Differential expression analysis between Front and Rear cells (**Fig. 2F**) identified collections of genes associated with cell motility (e.g.*Tagln*, *Acta2*, and *Cav1*) enriched in Front cells and genes associated with cell division (e.g.*H2afv*, *Cenpa* and *Stmn1)* for Rear cells. We displayed the expression level of single genes from (Fig. 2F) onto the UMAP representation and found that each gene had slightly different patterns, indicating additional heterogeneity of the programs at this resolution (**Fig. 2G**). Gene ontology (GO) analysis of biological processes using significantly differentially expressed genes supported the broad segregation of cell division (Rear) and motility (Front) genes in these two regions (**Fig. 2H**). Furthermore, using gene signatures for S and G2M phases of the cell cycle^12^, we observed that the Rear population exhibited significantly higher signature scores for S and G2M phases (**Fig. 2I and J**). Cluster 1 was especially enriched for cells in S or G2M phase relative to cluster 0 and 2 which were largely in G0/G1 phase (**Fig. 2K**). Finally, we validated these findings, using antibodies detecting the proteins encoded by genes discovered by ZipSeq. Staining for ACTA2 was broadly enriched near the front (**Fig. 2L**), tapering after peaking around 100 µm from the wound edge (**Fig. 2N)** whereas staining for STMN1 demonstrated enrichment **(Fig. 2M)** approximately 300 µm from the front (**Fig. 2N)** consistent with the regions we defined.

### Mapping Cortex vs. Medulla in a Live Lymph Node

We next applied ZipSeq to learn examine gene expression in entire mouse lymph nodes (LN), which have well-characterized spatial organization of cell types and gene-expression. We targeted two regions for ZipSeq discovery; the ‘Outer’ cortex extending from the tissue edge to the T-B margin and an ‘Inner’ region largely comprising the deep T cell zone and the medulla. We first stained live LN sections from an adult C57Bl/6 mouse with fluorescently labeled anti-CD3ε and anti-B220 antibodies to delineate these regions by widefield microscopy (**Fig. 3A-B**).

**Figure 3:**
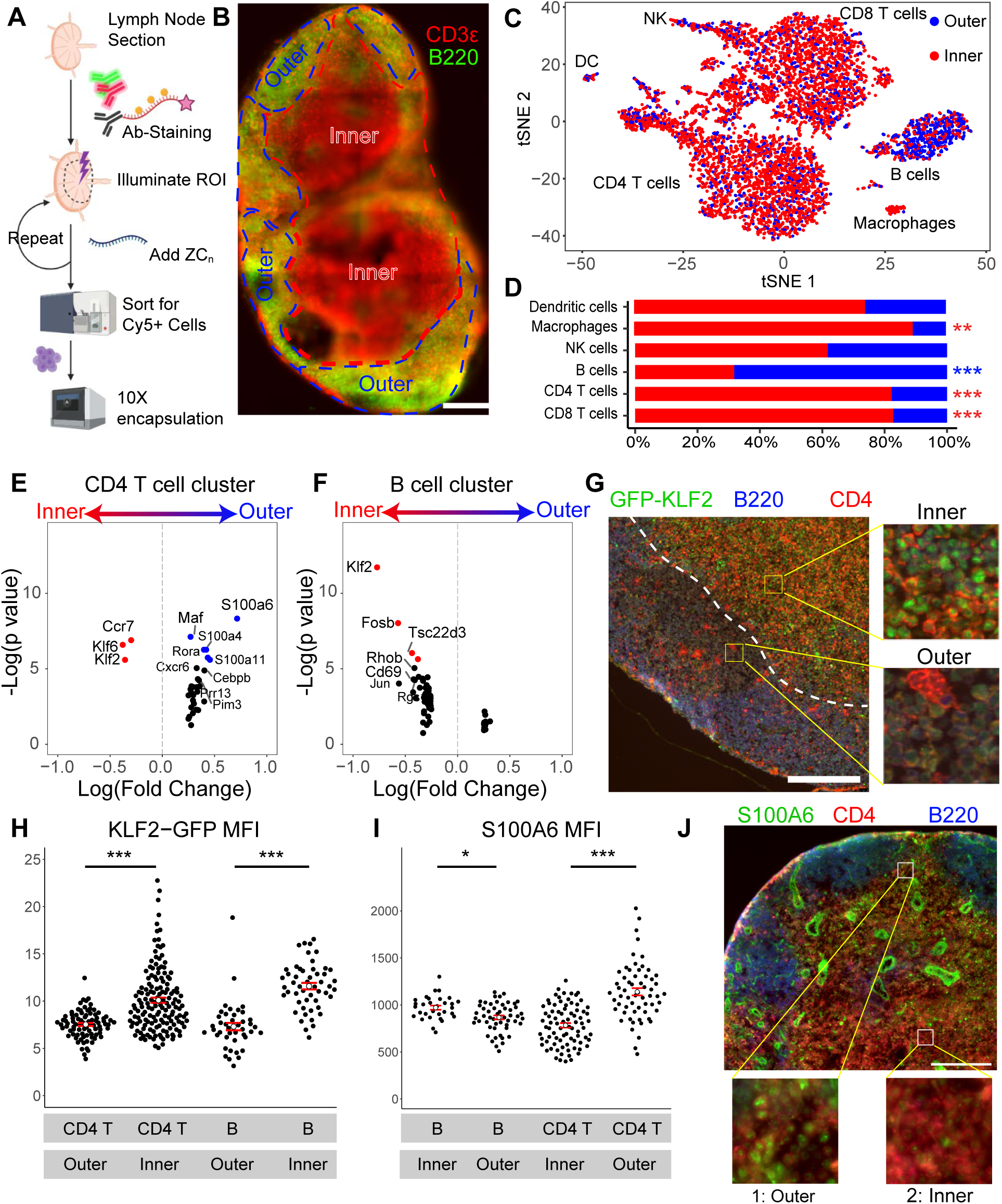
**(a)** Schematic of workflow for lymph node study. A lymph node was taken from a C57Bl/6 mouse and sectioned. Following this, the section was stained for B220 and CD3ε along with the anti-CD45 Ab conjugated oligo hybridized to a caged strand. The section was imaged and ROI’s were illuminated prior to Zipcode 1 or 2 addition. Tissue was then dissociated and labeled live cells (Cy5^+^) were sorted for 10X encapsulation. **(b)** Composite stitched image of lymph node section used with B220 marked in green and CD3ε in red to delineate inner and outer regions used for Zipcoding in subsequent study. Scale bar = 400 µm. **(c)** tSNE dimensional reduction of live cells. Assigned regional id based on ZC1:ZC2 counts overlaid. Immune cell populations were identified using known expression markers on Immgen. **(d)** Regional distributions of major immune cell populations as identified in (c). Asterisks denote significance of enrichment with color indicating direction (Inner vs. Outer). ***: p<0.0001 **: p<0.001 and *: p<0.01 **(e)** Volcano plot showing differential gene expression analysis within the CD4 T cell. Colored points represent genes with a p-value < 0.05 (Bonferroni adjusted). **(f)** Same as in e for the B cell population. **(g)** Immunofluorescence imaging of fixed lymph node section taken from a GFP-KLF2 reporter mouse. Section was stained for GFP, CD4 and B220. Dotted line represents demarcation between inner and outer regions used during quantification. Scale bar = 200 µm. Zoomed in insets show representative fields within inner and outer regions. **(h)** Mean fluorescence intensity of GFP-KLF2 signal intensity within CD4 T and B cells in IF image from (g) either inner or outer region. **(i)** Mean fluorescence intensity of S100A6 signal within CD4 T and B cells found in outer and inner regions of the lymph node in IF image from (j). Bee swarm plots represent intensities of individual cells with bars denoting standard error. *** p-value < 0.0001, ** p-value < 0.001 * p-value < 0.05 by Wilcoxon’s rank sum test. **(j)** Fixed frozen lymph node section stained for CD4, B220, and S100A6. Zoomed-in insets show representative fields from outer and inner regions, Scale bar = 200 µm.

We also labeled immune cells within the section with an anti-CD45 based caged anchor strand that was also conjugated to Cy5 to allow for purification of immune cells prior to encapsulation. Using the B220:CD3ε signal as a guide, we first printed Zipcodes to the Outer region and, following washing, printed a second Zipcode to the inner region. The lymph node section was then dissociated and live labeled CD45+ cells were sorted out based on Cy5+ signal and encapsulated. Following merging of Zipcode (ZC) and cDNA counts, we separated populations based on dominance of ZC1 and ZC2 (**Fig. S2**). In parallel, we performed tSNE dimensional reduction and identified major immune cell populations using compiled expression signatures on Immgen. We then overlaid ZC identity onto the tSNE projection. On first inspection, this revealed clear enrichment of “Outer” cells in the B cell cluster and “Inner” cells for T cell populations (**Fig. 3C and D**), consistent with expectations and fluorescence imaging.

Using the ZC spatial information, we performed differential gene expression analysis within populations based on position. Within the CD4 T cell population, we identified *Ccr7*, *Klf2 and Klf6* preferentially expressed in cells found in the Inner region and calcium binding proteins *S100a6/4* and transcription factor *Rora* preferentially expressed in the Outer region (**Fig. 3E**). Performing a similar analysis in B cells identified *Klf2* and *Fosb* in B cells found in the Inner vs. Outer region (**Fig. 3F**). Given the appearance of *Klf2* as an “inner” enriched gene in both of these analyses, we validated its spatial expression pattern using immunofluorescent imaging of lymph node sections **(Fig. 3G)** from a previously generated KLF2-GFP reporter mouse^13^. Using B220 and CD4 staining to identify B and CD4 T cells respectively, we found that there was indeed more KLF2-GFP expression in both B and CD4 T cells in the interior (**Fig. 3H**). We similarly validated that T cells found near and in B cell follicles expressed more S100A6 than those found deeper in the T cell zone (**Fig. 3I and J**). We noted a more modest difference in S100A6 in B cells (**Fig. 3I**). We also observed this particular protein in cells comprising high endothelial venules (HEV), which had been excluded from Zipcode analysis due to our selection of an anti-CD45-based anchor.

### Mapping immune cell differentiation in relation to position within tumors

To map variations in immune cell expression state within a live tumor, we derived cell lines from spontaneous tumors arising in the PyMT chOVA mouse breast cancer model, in which mCherry and ovalbumin (OVA) were co-expressed under the MMTV promoter, along with the Polyoma middle T antigen (PyMT)^14^. We orthotopically injected these into the inguinal mammary fat pad of female C57Bl/6 mice and 10 days later, we adoptively transferred 2 million CD8 T cells isolated from an OT-I UBC-GFP mouse. We allowed the T cells to expand for 4 days in lymph nodes and traffic to tumors. We then harvested the tumors and sectioned them into ∼150 µm thick slices (**Fig. 4A**) and observed dense clusters of GFP OTI T cells in the tumor margin with more dispersed cells in the interior (**Fig. 4B**).

**Figure 4:**
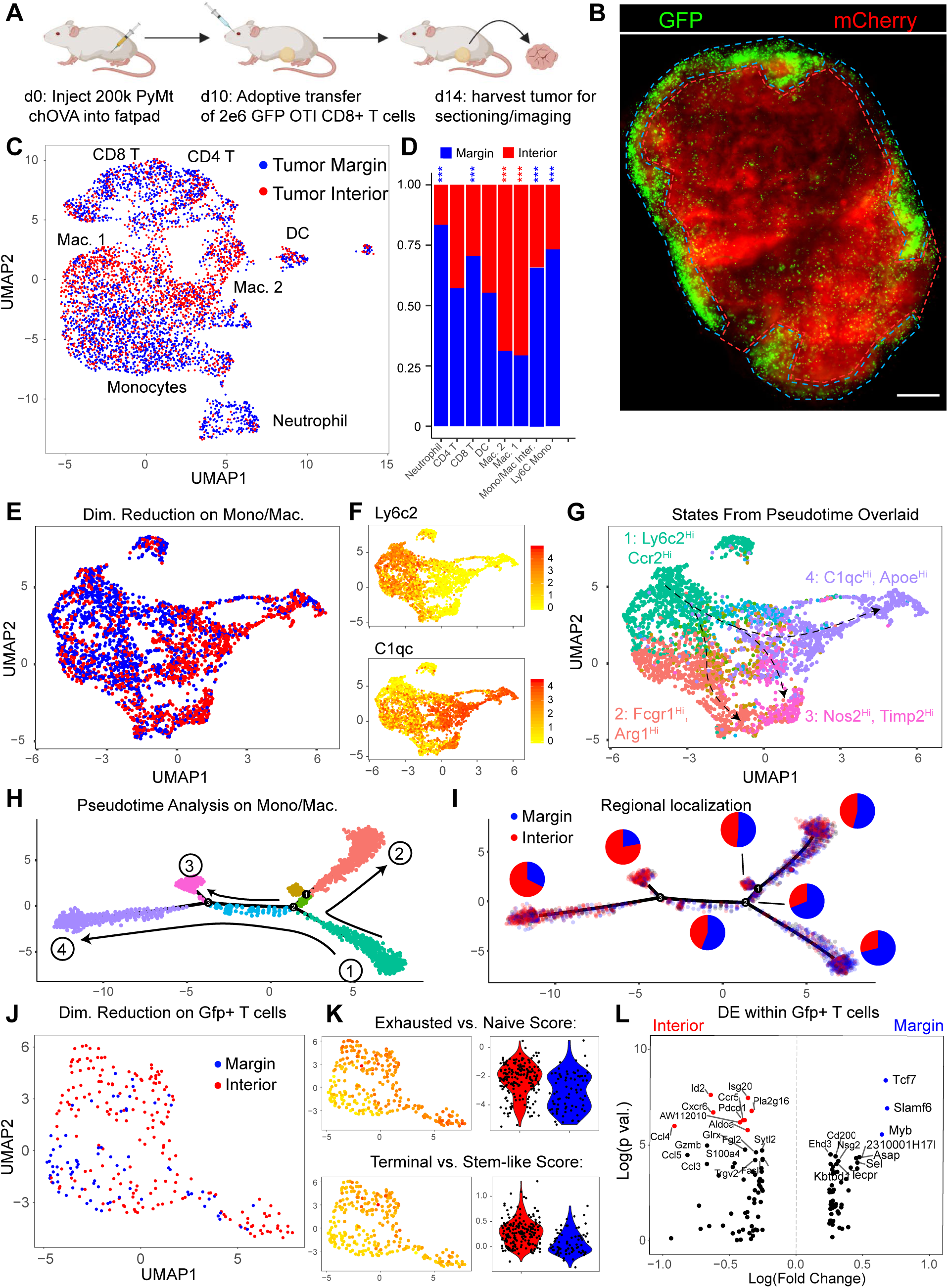
**(a)** Schematic of experimental setup. 200k PyMT-ChOVA tumor cells were injected into the inguinal mammary fatpad of 8-week-old female C57Bl/6 mice. After 14 days, 2e6 CD8 T cells from a GFP OTI mouse were adoptively transferred. Following 4 more days, tumor was harvested, sectioned, imaged, and labelled with Zipcodes according to the ROI’s denoted in **(b).** Imaging of 180 µm live tumor section used for scRNA-Seq in following experiments. Red channel denotes mCherry signal from PyMT-ChOVA tumor cells and green channel denotes adoptively transferred GFP OTI T cells. ROI’s used for Zipcode labeling shown overlaid. Scale bar = 400 µm. **(c)** UMAP representation of live labeled CD45+ cells. Cells below nUMI and ZC count threshold or above mitochondrial percentage threshold were filtered out. Assigned regional id based on ZC1:ZC2 counts overlaid. Large scale populations annotated based on similarity to known markers on Immgen. **(d)** Stacked bar charts denoting regional distributions of major immune cell populations from data in (c). **(e)** UMAP dimensional reduction on Monocyte/Macrophage population subset with regional identity as determined by ZC1:ZC2 ratio. **(f)** Feature plot of UMAP representation in (e) with normalized gene expression denoted by color scale for *Ly6c2* as a marker for monocytes and *C1qc* as tumor-associated macrophage (TAM) marker. **(g)** UMAP representation of monocyte/macrophage population with state identity calculated from Monocle pseudotime analysis in (h) overlaid. Arrows represent differentiation trajectory from the monocyte population to the terminal macrophage populations. Each major state is annotated with a selected marker genes. **(h)** DDR Tree dimensional reduction of monocyte/macrophage population as computed by Monocle with state identities overlaid. Arrows denote increasing pseudotime with the *Ly6C*^Hi^, *Ccr2*^Hi^ state designated as the root. **(i)** DDR Tree dimensional reduction of monocyte/macrophage population plotted with regional localization overlaid. Pie charts represent regional distributions (marginal vs. interior) for each state. **(j)** UMAP dimensional reduction on cells within the T cell clusters expressing at least one GFP transcript. Regional identity as determined by ZC1:ZC2 ratio overlaid. **(k)** UMAP representation with gene expression signature scores overlaid. Exhaustion vs. naïve gene signature scores were calculated for the *Gfp*+ T cell subpopulation (cells within lymphoid clusters with at least one *Gfp* transcript) and these scores overlaid on the UMAP representation. Violin plot represents this score distribution based on regional identity. Bottom row represent similar quantification of a terminal vs. stem-like exhausted signature score. **(l)** Volcano plot showing top differentially expressed genes in the *Gfp*+ T cell subpopulation based on regional identity. Colored points represent genes with a p-value < 0.05 (Bonferroni adjusted).

We defined “margin” vs. “interior” regions based on the GFP and Cherry signals and used anti-CD45-Cy5 labeled anchor strands and light-based to label them with Zipcodes 1 and 2, respectively (**Fig. 4B**). After dissociation of the tumors into a single cell suspension, we sorted out Cy5+ cells and encapsulated them for scRNA-Seq using our modified 10X scRNA-Seq workflow. Dimensional reduction and nearest neighbor clustering on RNA levels from these cells revealed clusters of T lymphocytes and monocytes/macrophages (**Fig. 4C**) and smaller populations of neutrophils, dendritic cells, and NK cells. Several of these populations displayed distinct regional distributions within the sample. For example, lymphocytes and neutrophils skewed towards the marginal region while macrophage populations were more frequently in the interior, matching observations made in a subcutaneous colorectal cancer tumor model using whole-volume imaging and tissue clearing^15^ (**Fig. 4D**) Sub-sampling only the monocyte/macrophage cluster (**Fig. 4E**) we found that that “margin” cells appeared differentially enriched on the left hand side of the UMAP projection where prototypical monocyte genes like *Ly6c2* were also enriched. Conversely “interior” cells were predominantly found on the right hand side where terminal tumor-associated macrophage (TAM) markers such as *C1qc and Apoe* were enriched (**Fig. 4E and F**)^16^. Exploring this further, we generated a pseudotime trajectory using Monocle^17^ with the *Ly6c*^Hi^, *Ccr2*^Hi^ state as the root state (State 1) (**Fig. 4H).** We observed differentiation of several TAM states (States 2-4) from the root monocyte state as pseudotime advanced with graded changes in gene expression such as loss of *Ly6c2* and *Ccr2* expression and gain of other TAM defining markers (**Fig. S3**). These terminal TAM states could be defined by expression of marker genes (**Fig. 4G and Fig. S4**) consistent with previously described TAM markers^18^. When we overlaid regional localization onto our pseudotime trajectory, we observed the regional localization of cells shift from margin to interior as pseudotime progressed from State 1 to terminal states 2,3 and 4 (**Fig.4I**). The terminal states exhibited their own differences in regional localization with State 2 more marginal vs. States 3/4 (**Fig.4I**) and differentially expressed genes based on localization (**Fig.S5**). One common feature amongst each ‘arm’ was an enrichment for monocyte markers *Ly6c2*, *Ccr2* and *Plac8* in marginal cells further supporting a gradient of differentiation concurrent with infiltration depth^19^.

Focusing on the antigen-specific (*Gfp*-expressing) OTI T cells, we also observed regional variation in the distribution of recently arrived effector CD8 T cells in transcriptional space (**Fig. 4J**). We observed a clear enrichment for genes previously associated with exhaustion (vs. naïve) in interior localized antigen specific T cells (**Fig. 4K**)^20, 21^. Given that T cell exhaustion represents a graded process, we also applied a terminal vs. stem-like exhaustion signature and observed a clear increase in terminal vs. stem-like exhaustion score for interior cells (**Fig. 4K**)^22^. Similarly, when we performed differential expression analysis for marginal vs interior cells in this GFP+ subset, the most significant gene hits were enriched in those defining earlier differentiation (e.g. *Tcf1* and *Slamf6*) in marginating T cells and more committed exhaustion (e.g. *Id2* and *Pdcd1)* in the interior T cells **(Fig. 4L**).

### Increased resolution in lymph node reveals spatial patterns of gene expression

To increase the number of regions that could be labeled, we devised two variations of ZipSeq. Instead of a terminating Zipcode sequence, we used DNA duplex strand bearing an orthogonal NPOM caged overhang (O2) sequence to effectively swap the potential binding site from O1 to O2 upon illumination in the first round. Thus, multiple regions can be defined at a time, through downstream addition of Zipcodes bearing either a complement to overhangs O1 or O2. This approach can theoretically be scaled up through the use of additional orthogonal overhang sequences, resulting in definition of 2^N^ regions using N+1 rounds of illumination and oligonucleotide addition. We demonstrate our ability to print a series of strands using two fluorescently labeled oligonucleotides as seen in **Fig. 5A** to generate 4 distinct regions.

**Figure 5:**
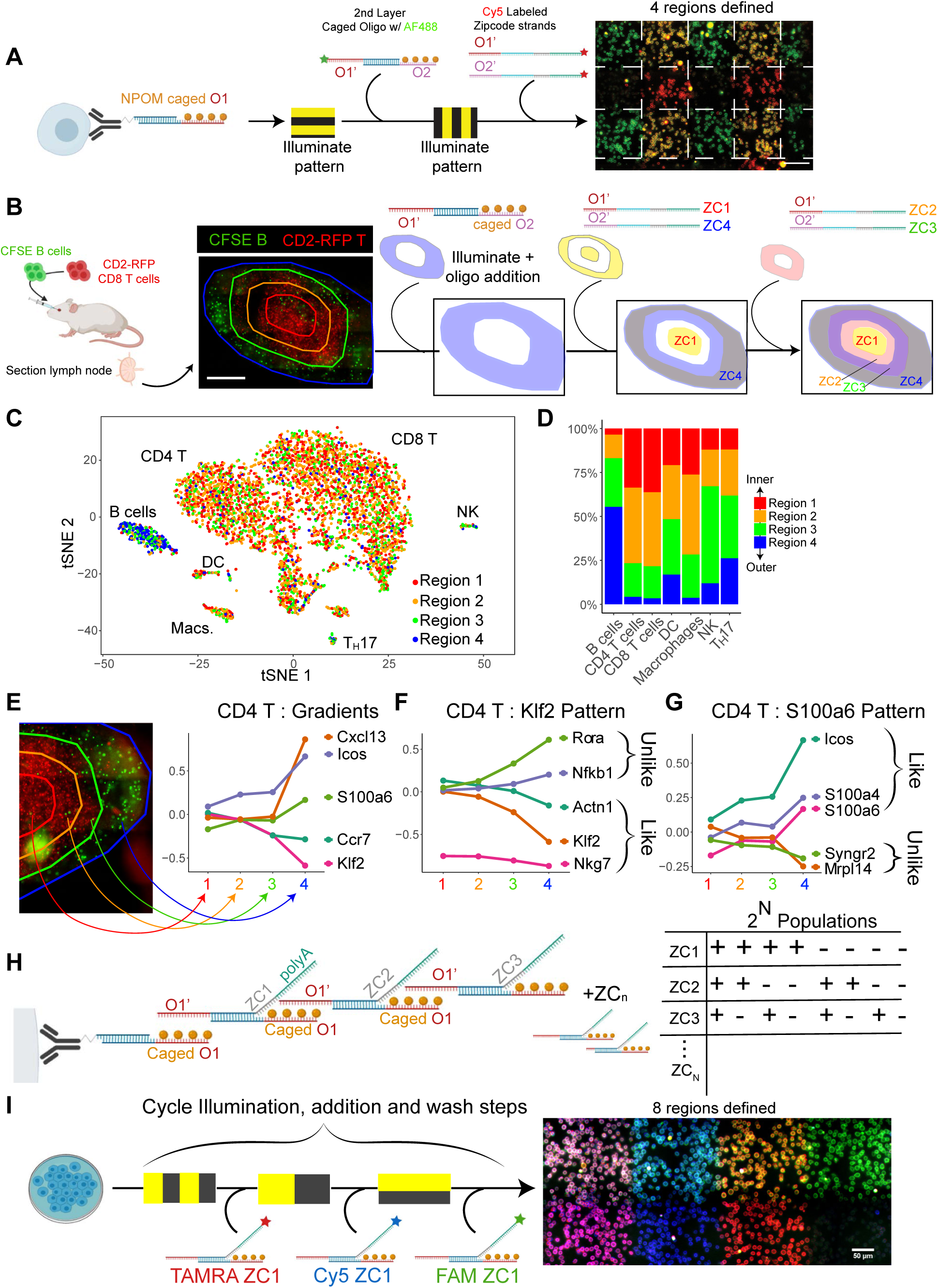
**(a)** Schematic of oligonucleotide design used for defining 4 regions by adding a second layer of caged oligonucleotides. A secondary oligonucleotide duplex bearing an orthogonal caged O2 overhang can hybridize to the uncaged O1 sequence. With a combination of Zipcode strands with either an overhang region O1’ or O2’, 4 distinct regions can be defined by 4 Zipcode species. Workflow illustrates the ability to define 4 such regions through fluorescence tagging of these oligonucleotide strands. Scale bar = 100 µm. **(b)** B cells extracted from C57Bl/6 mice were labeled with CFSE and CD8 T cells were extracted from a CD2-RFP mouse and adoptively transferred into a C57Bl/6 mouse. Lymph nodes were harvested, sectioned and imaged. 4 regions were defined with a unique Zipcode1-4 as overlaid onto the micrograph. These 4 regions were generated using the sequence of illumination and oligonucleotide additions shown. Scale bar = 400 µm. **(c)** tSNE dimensional reduction of live, labeled cells from LN section shown in (b) with regional identity overlaid. Major immune populations are annotated. **(d)** Bar chart illustrating distribution of cells in each of the 4 regions for selected immune cell populations in (c)**. (e)** Plots of mean scaled gene expression levels within the CD4 T cell cluster as a function of regional assignment for selected genes. **(f)** Plots of average scaled gene expression levels within the CD4 T cell cluster as a function of regional assignment for *Klf2* and two similar and dissimilar genes as calculated by cross-correlation score. Genes with significantly different expression levels and a logFC threshold of 0.4 between at least one pair of regions were considered for analysis. Cross-correlation scores were calculated between the averaged scaled expression levels of these genes and the reference gene. **(g)** Similar analysis with *S100a6* as reference. **(h)** Schematic of second design iteration. An anchoring moiety is conjugated to an anchor sequence which is hybridized to an oligonucleotide with a NPOM caged overhang sequence. Each additive Zipcode duplex block consists of a Zipcode strand with a complement to the overhang sequence, a universal hybridization sequence, an 8 nt barcode, and a 28 nt polyA sequence. Meanwhile, a strand with a caged overhang is hybridized to the Zipcode sequence through the universal hybridization sequence. In this way, Zipcode blocks are added on in a combinatorial manner, defining 2^N^ populations based on presence or absence of a given Zipcode block. **(i)** Demonstration of combinatorial spatial barcoding of a field of cells in an exponentially scaling manner. Conjugate labeled CD8 T cell were plated and subjected to a 3X sequence of illumination patterns and Zipcode block additions bearing distinct fluorophores resulting in 8 regions with distinct fluorophore combinations. Scale bar = 50 µm.

Using this design, we sought to test our method for its ability to detect gradients of genes, such as those for chemokine receptors like CCR7, in the lymph node. We adoptively transferred B cells (CFSE-labeled) and CD8 T cells (RFP-labeled) into a C57Bl/6 mouse before harvesting and sectioning the inguinal LN which revealed T and B zones (**Fig.5B**). The sequence of illumination patterns and oligonucleotide additions to generate 4 concentric regions of labels (ZC1-4) is illustrated in **Fig.5B**. Following dissociation and sequencing, we observed 4 groupings of cells with dominant ZC counts for 1-4 while filtering out cells that were ambiguous with no clear ZC dominance(**Fig. S6**). When we overlaid these regional identities onto our tSNE representation, we observed a strong enrichment for regions 3,4 in the B cell cluster vs T cell clusters (**Fig.5C**). We noted that the degree of region 4 enrichment in B cells vs. T cells (>50% vs. 4%) (**Fig.5D**) was greatly increased compared to our 2-region lymph node study in **Figure 3**. In addition, when examining rarer immune cell subpopulations, we observed the natural killer cell population enriched specifically at the interface between T and B cell zones (region 3) as has been previously observed^23^. (**Fig.5D**).

Definition of 4 concentric regions should permit us to identify the existence of gradients of gene expression in space. As predicted, within T cell clusters, expression of *Ccr7* and *Klf2* steadily decrease moving outwards towards the B cell zone while *S100a6* expression increases, matching our findings from Figure 3 and published results^24^(**Fig. 5E**). In contrast, *Cxcl13*, a chemokine very highly associated with B cells zones, was significantly expressed in region 4 alone (**Fig. 5E**). This type of analysis allowed us to compare patterns of gene expression across space using genes such as *Klf2* and *S100a6* as references, and calculating cross correlation scores with all other expressed genes in our dataset, identifying the most ‘similar’ and ‘dissimilar’ genes. For example, *Klf2* shares similar profiles with *Nkg7* and *Actn1* (with which it shares a similar differentiation trajectory in vitro)^25^ while being most dissimilar to *Rora* and *Nfkb1*. Meanwhile *S100a6* shares similarity to its frequent binding partner *S100a4* as well as the costimulatory molecule *Icos* while being most dissimilar to *Syngr2* and *Mrpl14* (**Fig. 5F,G**).

This approach, while allowing for definition of exponentially increasing number of regions, also requires a similar scaling of distinct orthogonal caged sequences which could become cost-prohibitive, so we devised a second variation. Here, each coding segment consists of a Zipcode ‘block’ which is a duplex of a polyA, barcode, a universal hybridization region, and an overhang sequence O. This strand is pre-hybridized to a strand with the universal hybridization region and a caged overhang sequence O’ (**Fig. 5H**). This schematic necessitates synthesis of only a single caged sequence species and N distinct barcoded polyA strands yielding potentially 2^N^ regions after N rounds of illumination and addition. Using an in-tube validation experiment, we were able to observe 4 separate populations of cells using two rounds of illumination and Zipcode addition (**Fig. S7**). Merging of cell type identity and Zipcode combination showed good agreement with the experimental scheme. To spatially visualize this approach, we used 3 separate illumination and addition steps with three distinct Zipcode blocks, each bearing a distinct fluorophore. This yielded 8 (2^3^) distinct color combinations, each defining a grid position (**Fig. 5I**). Because the resolution of this method is theoretically diffraction limited, we repeated this using smaller regions, approximately 20 µm per side, demonstrating the capability to define areas on the order of single cells or small neighborhoods (**Fig. S8**).

## DISCUSSION

Here we introduce ZipSeq, an approach that allows for on-demand barcoding of cells within defined regions during microscopy. We demonstrate that our approach is compatible with live tissue sections and with multiple cell and tissue types, depending on anchoring moiety. It precludes the need for genetically encoded photoactivatable proteins^26^ so is applicable to human tissues and allows for definition of multiple regions at once. ZipSeq plugs into the commercially available 10X workflow^27^, and is theoretically compatible with many other scRNA-Seq methodologies^28–30^ making its wider adoption feasible, requiring only caged oligonucleotides and a photo-patterning module.

Using this approach, we demonstrate the ability in both an in vitro wound healing model and ex-vivo tissue sections (lymph node and tumor) to assign single cell transcriptomes to regions defined concurrently with fluorescence imaging. In the wound healing model, our approach identified distinct transcriptional programs activated in fibroblasts as a function of distance from the wound edge with several targets validated through immunofluorescence. We found a migratory cell state enriched at the leading edge (0-200 µm) and a proliferative state enriched at the rear (200-400 µm). The generalized spatial segregation between migration and proliferation has been previously observed in multiple cell types such as epithelial cells and keratinocytes during wound healing^31, 32^ with overexpression of *Acta2* and *Serpine1* described at the leading edge in prior wound healing studies^33, 34^. It will be informative to repeat this study at higher resolution at different timepoints to observe how these spatial patterns of expression might evolve or whether patterns exist in other axes or at other scales.

In the context of lymph nodes, this method reports spatially dependent gene expression validated by previous works including *Klf2*, *Ccr7* and *S100a6* expression^13, 35, 36^. With increasing region number, this method permits identification of genes that map ‘similarly’ or ‘dissimilarly’ to a known gene over space, some of which are known, while others are not. Additionally, in the context of a tumor model, this method allows the progression of myeloid and T cell differentiation to be mapped to physical infiltration depth. The myeloid differentiation in particular is consistent with recruited monocytes receiving local cues that skew differentiation trajectories as they arrive in the tumor margins, as described previously in several tumor models^37, 38^.

Alongside macrophage differentiation, we also observed genes associated with T cell exhaustion upregulated in tumor specific CD8 T cells when comparing tumor interior with margins. Notably *Tcf1*, a major factor in maintenance of a stem-like exhausted phenotype, was enriched in marginal T cells vs. interior^22, 39^. Comparison with imaging data taken prior to barcoding suggests that the dispersed, deeper infiltrating antigen-specific T cells we observed (**Fig. 5B**) are further along the exhaustion pathway compared to the T cells at the edges. The mechanistic link between depth and commitment towards exhaustion bears further investigation. We also noted the enhanced expression of chemokines/receptors in interior vs. margin CD8 T cells associated with increased trafficking and infiltration of T cells into the tumor such as *Ccl4/5* and receptor *Ccr5* (**Fig. 4L**)^40–42^.

Many spatial transcriptomics approaches use prefabricated grids of barcoded poly-dT’s or barcoded beads bearing poly-dT’s to capture all transcripts within the grid position^5–7^. While these approaches offer excellent spatial resolution, it will be difficult to apply them directly to tissue following live imaging or perturbations. The relatively low read depth offered by several of these approaches (∼1e4 reads per 150 µm grid square for Spatial Transcriptomics and 2e2 per 10 µm bead for SLIDE-Seq), would further complicate analysis of expression patterns in rarer cell types or those with low levels of RNA as their transcripts become diluted out during the capture step. Because ZipSeq plugs into droplet-based scRNA-Seq workflows, it has potential to tap into greater read depth per cell, generating true single cell transcriptomes without the need for deconvolution of a pool of transcripts derived from multiple cells. Another advantage of ZipSeq is the potential to easily integrate with other multimodal measurements such as concurrent surface epitope labeling using CITE-Seq, ATAC-seq or single cell immune repertoire sequencing^8^.

ZipSeq however faces limitations, most notably in spatial resolution. We propose that ZipSeq is currently most effective in questions based around microanatomical features observed during imaging that can guide definition of ROI’s for barcoding. To make a layered barcoding printing scheme achievable and cost-effective, we demonstrated the ability to add on layers of secondary caged oligonucleotides to increase the number of definable regions. With this increased spatial resolution, we can describe the segregation of different cell states in finer detail and detect genes with sharply defined spatial expression in an unbiased fashion.

While many of our studies focused on the immune compartment, alteration of anchoring moiety will potentially expand application to diverse multicellular models. Combined with live imaging of cells prior to Zipcoding, time-dependent cell behaviors such as motility can be used to define regions of interest (e.g. transcriptional states of cells in low and high motility zones, zones defined with measures of hypoxia, etc.). In summary, ZipSeq represents a novel approach to mapping scRNA-Seq data from conventional scRNA-Seq workflows using on-demand light-controlled hybridization of DNA barcodes onto cells. We propose that ZipSeq will strengthen our capability to link spatial heterogeneity in multicellular systems to transcriptional heterogeneity of the constituent cell populations.

## ACKNOWLEDGEMENTS

We thank the Biological Imaging Development Center at UCSF for help with microscopy data collection and instrumentation. We also thank the Parnassus Flow Cytometry Core for flow cytometry instrumentation and the Institute for Human Genomics at UCSF for sequencing and bioinformatics support. In addition, we thank the Computational Biology and Informatics (CBI) core at the UCSF Helen Diller Family Comprehensive Cancer Center (HDFCCC) for computing resources. This work was supported from NIH/NCI grants P30DK063720 and 1R01CA197363, by the Parker Institute for Cancer Immunotherapy (PICI) as well as NIH T32 training grant (5T32CA177555-02)

## AUTHOR CONTRIBUTIONS

K.H.H. conceived of the experiments, performed the experiments, analyzed data, and wrote the manuscript. M.F.K. conceived of experiments, provided administrative and financial support, and wrote the manuscript. K.K. generated the PyMT chOVA cell line. C.S.M., D.S.P., E.D.C., and Z.J.G. generated the lipid anchored oligonucleotide. J.E. developed custom interface for controlling illumination patterns during imaging. S.J. generated the KLF2-GFP reporter mouse and associated IF data. A.A.R. assisted with analysis of scRNA-Seq data.

## Methods

### Oligonucleotide list

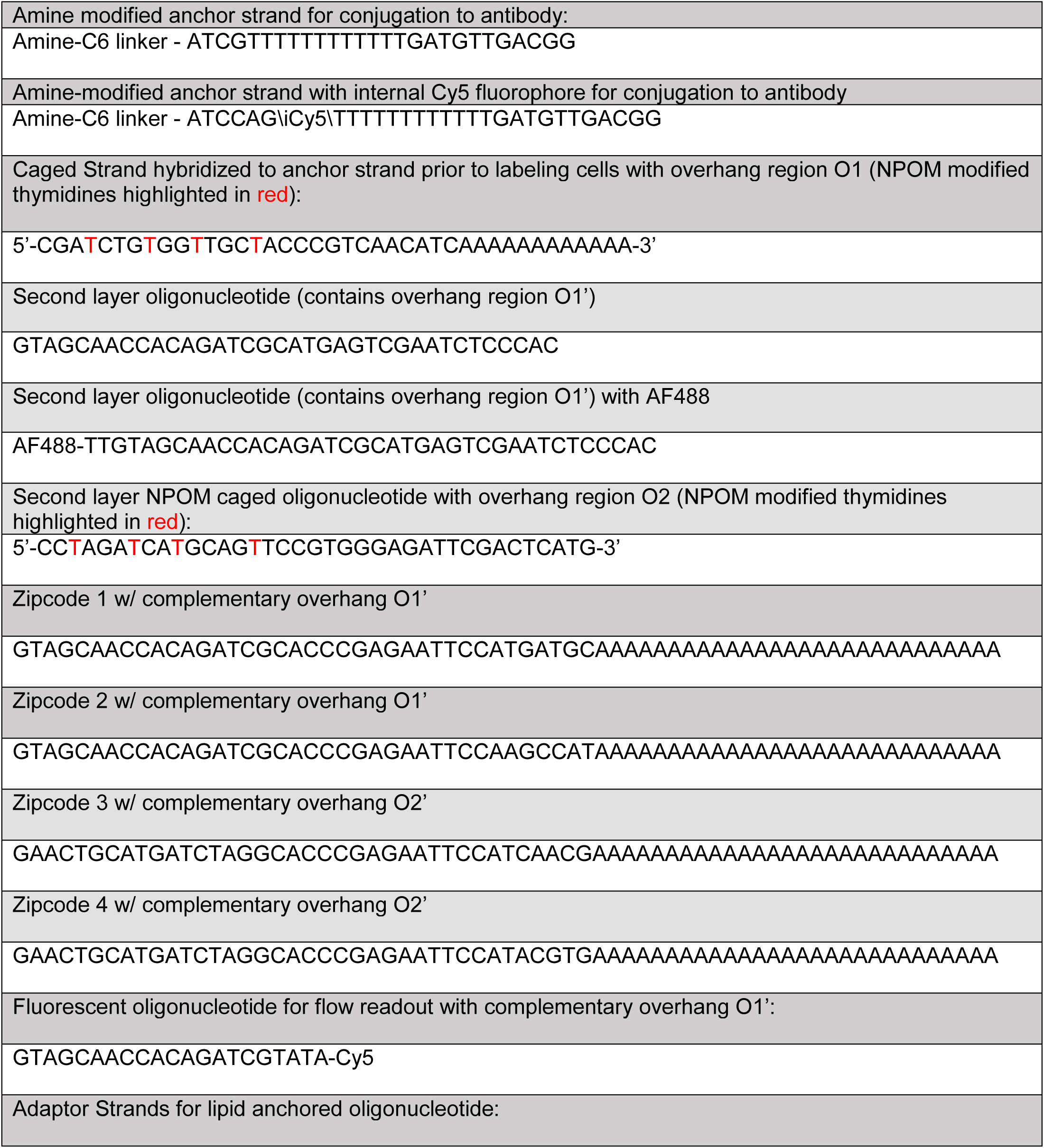
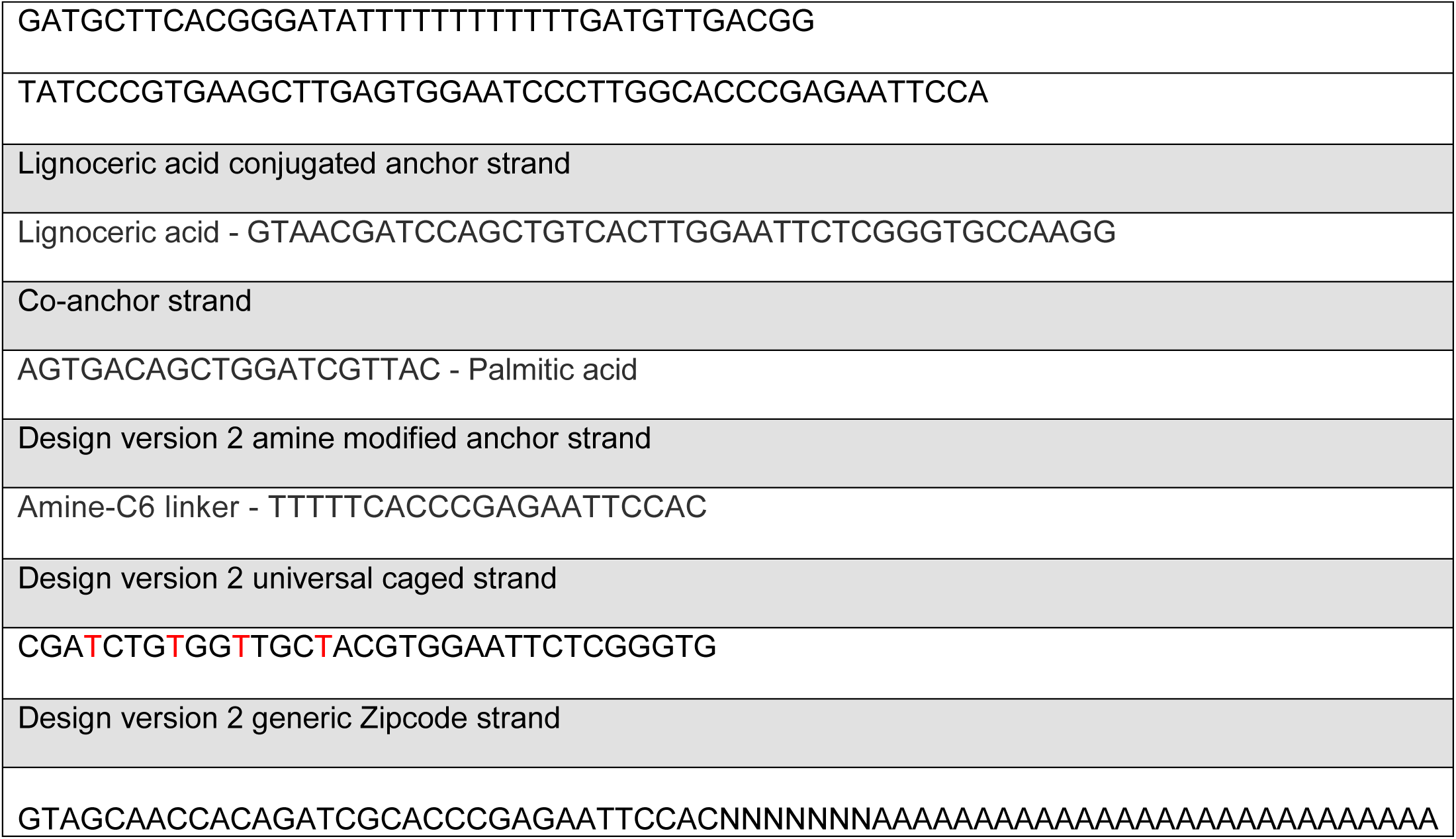

All oligonucleotides save for the caged strand were ordered from IDT with HPLC purification. NPOM caged strand was ordered as a custom synthesis from BioSynthesis.

#### Reagents

Nuclease Free Bovine Serum Albumin purchased from VWR

Single stranded salmon sperm DNA purchased from Life Technologies

10X v2 and v3 kits were purchased from 10X Genomics. SPRI selection beads from Beckman Coulter. Collagenase I and IV were purchased from Worthington Biosciences. 2X Kapa HiFi HotStart Master Mix.

RPMI 1640 and DMEM were ordered from Gibco.

#### Antibodies

LEAF-purified anti-CD45 antibody (30F-11), PE-anti-CD3ε (145-2C11), and FITC-anti-CD45R(B220) (RA3-6B2) purchased from Biolegend.

For IF, Abs for targets included anti-STMN1 (Abcam 52630), anti-S100A6 (Invitrogen PA5-16590), anti-ACTA22 (Sigma 1A4)

### Conjugation of Anchor oligonucleotide w/ Antibody

The Thunder-Link Plus kit (Expedeon) was used to conjugate the amine modified anchor strand to an anti-mCD45 antibody (clone 30F-110) at a molar ratio of 1:5 Ab:oligo and allowed to conjugate overnight at room temperature prior to conjugate purification according to instructions.

#### Lipid conjugated oligonucleotide

Synthesis of the lipid conjugated anchor oligo was performed as in^11^. In order to hybridize the caged oligonucleotide species, two adaptor sequences were also pre-hybridized with the anchor and caged strands (sequences shown above)

#### Mouse Strains

Experiments were performed in 6-8 week old female C57Bl/6J mice from JAX (#000664). For adoptive transfer, CD8 T cells were derived from a CD2-dsRed mouse (MGI# 5296821) and a OTI (C57BL/6-Tg(TcraTcrb)1100Mjb/J) (#003831) crossed with a UBC-GFP (C57BL/6-Tg(UBC-GFP)30Scha/J) (#004353).4

#### In-tube validation of zipcoding

The caged strand was pre-hybridized to the anti-CD45 Ab-anchor conjugate by adding a 1:1 molar ratio of caged to anchor strand and incubating at 37C for 15 minutes and allowing to cool to RT. CD4 and CD8 T cells were isolated from a mouse using CD4 and CD8 negative selection kits respectively (StemCell Technologies) and were labelled with the caged strand hybridized to the Ab-oligo conjugate. In the first round, CD4+ T cells were illuminated with 365 nm light and the first Zipcode strand added to both populations. Following a 4 min. incubation and 3 washes, the CD8 population was then illuminated and Zipcode 2 added and allowed to hybridize. Following a series of washes, the cells were pooled and encapsulated using a 10X v2 3’ kit with a target cell # of 10k.

#### Microscopy

Imaging and photo-uncaging was performed with a customized Zeiss Axiovert 200M inverted microscope fitted with a Mosaic digital micromirror device from Andor.

Illumination for wide-field imaging was provided by a Lambda DG-4. For photo-uncaging, light from a mercury arc lamp was passed through a 365 nm bandpass filter then directed into the DMD. Imaging was performed with a 20X objective from Zeiss (Plan-NEOFLUAR). For imaging, we used a Photometrics Evolve 512. Metamorph (Molecular Devices) was used to control the microscope and illumination by the Mosaic DMD with a custom Visual Basics program to provide a user interface for manual ROI delineation across multiple fields of view.

#### On-Scope validation of spatially controlled DNA hybridization

Isolated primary mouse CD4 T cells were plated in a Labtek chamber slide coated with anti-CD3ε antibody to adhere cells. Cells were then labeled with Ab-oligo conjugate hybridized to the caged strand. Following 2x washes and blocking w/ ssDNA to prevent nonspecific DNA-well interactions, vertical bands of 200 µm width were illuminated with 365 nm light from a mercury arc-lamp through the Mosaic. A fluorescently labeled Zipcode strand was flowed into the well and incubated for 5 minutes for hybridization. Following 3x washes with media, the sequence was repeated 2x for 2 other fluorescently labled Zipcode strands. The illuminated regions were then imaged.

For the combinatorial design shown in Fig. 5H and Fig. S11, Zipcode strands 1-3 were conjugated to fluorophores Cy5, TAMRA and FAM. These Zipcode strands were then hybridized to the complementary caged oligonucleotide strand to form 3 Zipcode blocks. Similar to the workflow above, a series of illumination, Zipcode addition, washing, and blocking generated 3 patterns of Zipcode hybridization overlaid, creating 8 distinct color combinations. The grids used in Fig. 5h were designed to generate 200 µm × 200 µm squares. In Fig. S11, the grid squares were scaled down to push the limits of DMD resolution to roughly 20 µm × 20 µm squares.

### Wound Healing study

NIH/3T3 cells were ordered from ATCC (CRL-1658) and cultured in complete growth medium (DMEM+10%FCS+55mM BME+PSG). 2 days prior to labeling and scRNA-Seq, a monolayer of 40k cells was plated in a well of a 8-well LabTek chambered coverglass. Cells were allowed to settle and fill in the well until 12 hours prior to imaging and labeling.

A pipette tip was used to introduce a scratch around 0.8mm in width in the monolayer. Cells were washed once with fresh growth medium to remove debris or floater cells and incubated for a further 12 hours. Cells were then transferred to the microscope and the regions of interest we delineated. Growth medium was washed out and replaced with serum-free phenol red free DMEM with ssDNA to block nonspecific DNA binding in further steps. The anchor-caged strand lipid tag was added and allowed to sit for 10 minutes followed by the co-anchor strand. Following 3 washes, the desired ROI’s were illuminated using a 800 ms pulse of 365nm light.

The first Zipcode strand was added to the monolayer and allowed to hybridize for 5 minutes. Following 2 washes, a blocking strand was added at a lower concentration for 5 minutes to prevent residual Zipcode strand from binding in the following cycle. After 2 another 2 washes, the second ROI was illuminated and the steps repeated with the second Zipcode strand.

After the last series of washes, the medium was removed and Accutase was added to detach the cells. After 5minutes of incubation at RT, cells were harvested and washed with cold PBS+0.04%BSA as recommended by 10X protocol. Encapsulation was performed for a target cell # of 10k using the v2 chemistry.

### Lymph node study

Inguinal lymph nodes were harvested from 8 week old C57Bl/6 female mice and embedded while live in 2% agarose. Using a Leica Vibratome, the lymph nodes were sectioned into 150 um thick slices and affixed to a LabTek chamber slide using Vetbond (3M) applied to the agarose ‘rim’. Sections were then incubated with anti-B220 FITC and anti-CD3ε PE along with the anti-CD45 conjugated NPOM caged anchor strand bearing an internal Cy5 modification for 1 hour at 4C. Following 3 washes, the sections were imaged on the scope and ROI’s delineated and illuminated. The first Zipcode was added at a 1 µM concentration in RPMI and allowed to incubate 10 minutes. Following 3 washes, the blocking strand was added at 0.25 µM concentration and incubated a further 5 minutes. Following a series of 3 washes, the process was repeated for the second Zipcode. Following a final blocking step, the section was mechanically disrupted and strained over a 100 µm nylon strainer and sorted for live, Cy5+ cells. Sorted cells were washed with cold PBS+0.04%BSA and encapsulated following 10X guidelines using a v2 3’ kit.

For the 4-region lymph node study, naïve CD8 T cells were purified from a hCD2-dsRed mouse and 4e6 were adoptively transferred into a 6 week old B6 mouse. Meanwhile, B cells were purified from a B6 mouse using an EasySep B cell negative selection kit (StemCell Technologies) and labelled with CFSE before being transferred at 4e6 per mouse. 3 days following transfers, the mouse was sacrificed and the inguinal lymph nodes extracted for sectioning and study.

### Tumor study

For orthotopically injected PyMTChOVA models, the PyMTChOVA breast cancer cell line was generated from *de novo* mammary tumors in the PyMTChOVA breast cancer mouse model^14^. Briefly, mammary tumors were harvested in ice cold PBS and mechanically minced into small fragments. Cells and tissue fragments were cultured in DMEM + 10% FCS and added Penicillin, Streptomycin and Glutamine. After 7-10 days tissue fragments and debris were washed out with ice cold PBS and attached cells were allowed to grow to confluency. Cells were cultured in growth medium for an additional 3-5 passages to generate the established PyMTChOVA mammary tumor cell line. Tumor cell line was kept frozen and thawed directly from stock as needed prior to injection to avoid unnecessary passages. 200k cells were injected in Matrigel (Corning) into the inguinal fat pad of 8-week-old female C57Bl/6 mice.

C57BL/6-Tg(UBC-GFP)30Scha/J mice were crossed to OT-1 mice to generate a GFP OT-1 mouse strain. Lymph nodes were harvested from a 6-week-old GFP OT-1 mouse and CD8 T cells were isolated using an EasySep mouse CD8 T cell negative selection kit (StemCell Technologies). 14 days post tumor injection, 5e6 CD8 T cells from a GFP OTI mouse were adoptively transferred through retro-orbital injection. 4 days following, the mouse was sacrificed, and the tumor harvested for sectioning on a Leica Vibratome into 150 µm thick slices. As before in the lymph node study, the tissue was embedded live into 2% agarose for sectioning. Sections were blocked with ssDNA and BSA in RPMI for 30’ at 4C then stained with the anti-CD45 Ab conjugated to NPOM caged anchor strand with internal Cy5 for 1 hr at 4C. Following washes, sections were affixed to an Ibidi µ-Slide 8 well using Vetbond. Sections were then imaged with the desired channels, and spatially Zipcoded as described above. Following the final block and wash step, tissue sections were diced finely and incubated with a Collagenase I and IV blend in RPMI and incubated for 30’ at RT. The resulting suspension was mechanically agitated by pipetting and then strained on a 100 µm strainer. As before, live Cy5+ cells were sorted out on a FACSAria II, washed in PBS + 0.04% BSA and then encapsulated following 10X specifications for v3 3’ chemistry.

#### Library Construction

cDNA library construction following GEM formation was performed as directed by 10X using v2/3 3’ chemistry depending on experiment. Following the published CITE-Seq protocol, an additive primer (partial Read 2 small RNA) was spiked into the cDNA amplification reaction. During the post-cDNA amplification SPRI cleanup step, the supernatant was saved and underwent two successive 3X SPRI cleanup steps. This fraction was expected to contain the shorter Zipcode reads. This library was then amplified using primers from CITE-Seq protocol^8^. Following fragment analysis on the BioAnalyzer and library quantification by qPCR, the Zipcode library was mixed with the associated cDNA library at a 1:10 molar ratio and sequenced on either Illumina HiSeq or NovaSeq using 10X recommended sequencing parameters.

#### Processing of raw sequencing reads

Raw read files were processed using CellRanger bcl2fastq to separate cDNA and ZC libraries. cDNA libraries were processed using standard CellRanger count function while ZC libraries were counted using a Python script made available by CITE-Seq using a whitelist of cells provided by the CellRanger count function determined by a minimum nUMI threshold^8^. The two outputs were then merged in Seurat for further analysis.

#### Data Analysis

Cells with a high mitochondrial read count % (above 10% assumed to be dead or dying cells) were filtered out. Cells with low counts for cDNA were filtered out based on the presence of a local peak at the low end of the distribution. Read counts were normalized using log-normalization, scaled and centered and nUMI and mitochondrial percentage regressed out. PCA was performed and the top 10 of these PC’s were used to inform the dimensional reduction by Seurat built-in tSNE or UMAP (using the Python implementation of umap-learn package). Meanwhile, nearest neighbor clustering using these PC’s was performed using Seurat’s built-in FindClusters function at a specified resolution of 0.8. In order to call Zipcode identities, Zipcode counts were normalized and the ratios of these normalized counts used to gate cells as belonging to one identity or another with ambiguous cells (between gates) filtered from analysis. For the 4-region lymph node study, k-means clustering on normalized Zipcode counts was used to generate 5 clusters corresponding to ZC1-4 dominance along with a centrally localized ambiguous population.

For differential gene expression analysis, we used Seurat’s built-in FindMarkers function which implements testing based on the Wilcoxon rank sum test along with a Bonferroni correction to adjust p-values. The testing was restricted to genes expressed in at least 10% of cells.

For Monocle analysis, we directly imported the Seurat object of interest into monocle and used the clusters pre-defined in Seurat to get a list of differentially expressed genes between clusters^17^. The top 800 genes were then used in the DDRTree dimensional reduction and pseudotime ordering.

For signature analysis, curated lists of genes from literature were passed into Single Cell Signature Explorer to generate scores^43^. Sources for gene lists used for signature scores derived from:

-S and G2M phase: from built-in gene lists in Seurat

-Exhaustion vs. Naïve:^21^

-Terminal vs. stem-like exhaustion:^22^

For analyzing spatial gradient profiles of genes in the 4-region lymph node study, we used the mean scaled and normalized gene expression for a given gene in each region. In order to find the most similar and dissimilar gene profiles, we first filtered out genes expressed in <10% of cells for all regions. We then calculated a cross-correlation score compared to a reference gene and ranked these scores from lowest correlation (−1) to highest correlation (1).

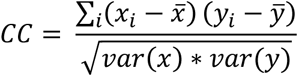

For gene x and y where i represents regions 1-4.

#### Immunofluorescence

For IF study on KLF2-GFP reporter mouse lymph nodes, sections were prepared, stained and imaged from PE-primed mouse on d14 as in^13^. For IF of S100a6, lymph nodes from 8 week old C57Bl/6 mice were embedded in optimal cutting temperature compound (OCT)(Sakura) and cryosectioned into 10 µm thick slices. Slices were fixed with 4% PFA and permeabilized by TritonX-100. Following blocking with 5% goat serum, primary antibodies were added and incubated at 4C overnight. Following washes, slices were then incubated with secondary Abs for 1 hr at RT, washed and then incubated for 5 minutes at RT with DAPI. Following another wash, slices were mounted with Vectashield and a coverslip, then imaged.

For IF studies on wound healing monolayers, NIH/3T3 fibroblasts were prepared in LabTek chambered slides as before and 12 hours post-wounding were fixed with 4% PFA, permeabilized with TritonX-100 and blocked with 5% goat serum.

Quantification of LN immunofluorescence: Analysis was performed within Fiji. Depending on the cell type of interest (CD4 T or B cells), CD4 or B220 signal was used to identify cells within the tissue. Identified cells were selected randomly based on this signal and MFI of target channel was calculated within this mask. Wilcoxon rank-sum test (two-tailed) used for comparison of MFI’s between populations.

For wound healing, binary masks for cell area were generated through thresholding of images and applied to target channel. Columns of 10 pixel width were taken progressively from wound edge and pixels in the mask were averaged to generate a column fluorescence intensity average.

### Statistical Testing

For quantification of immunofluorescence data, we performed a two tailed permutation test using the mean MFI as test-statistic. In order to see if regional enrichment for a given immune cell population from scRNA-Seq data was significant, we used a hypergeometric distribution to determine the probability of drawing a regional composition with equal to or greater enrichment at random. This probability was directly reported as the p-statistic.

---

**Supplementary Figure 1:**
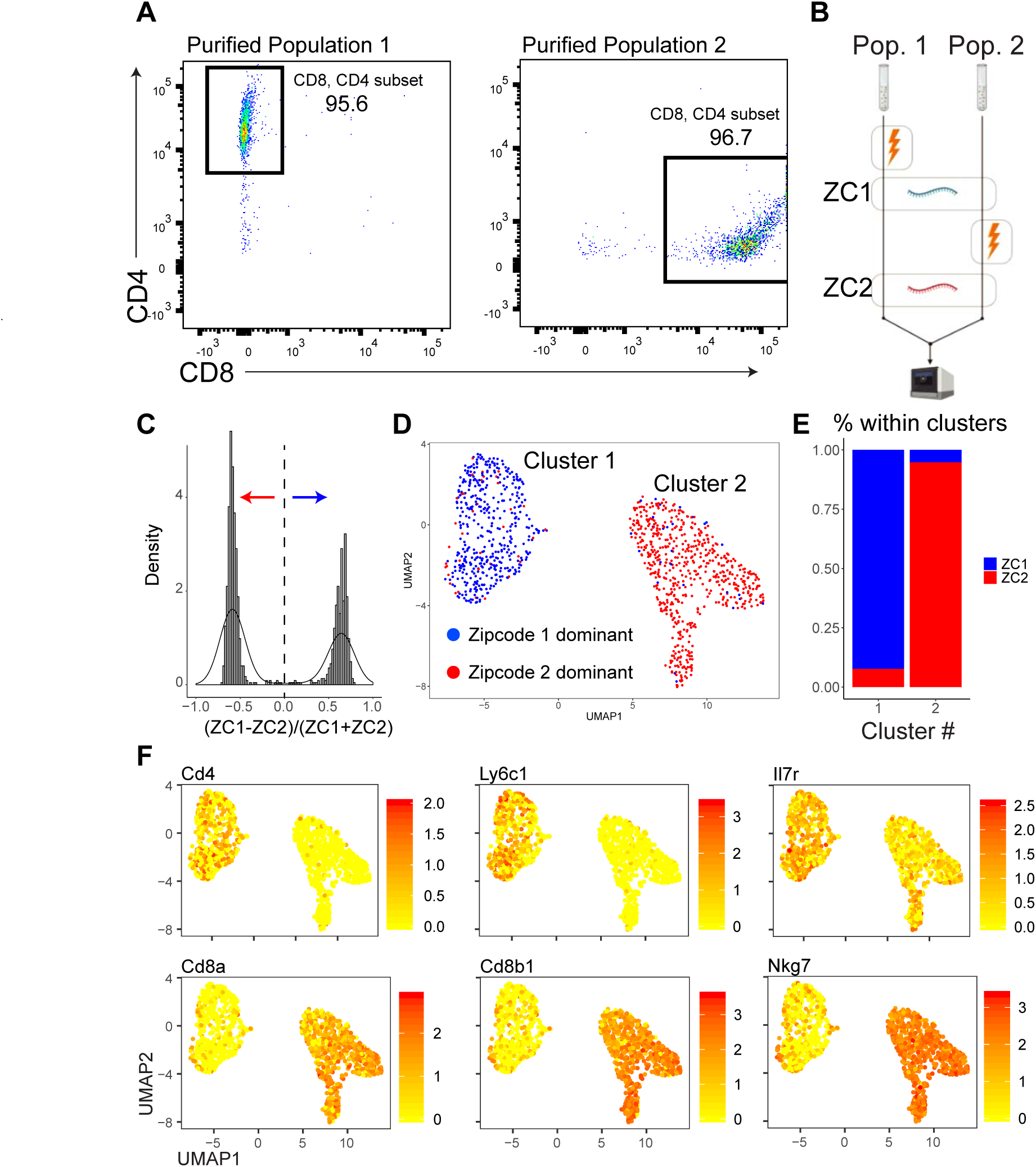
In-tube validation that photo-uncaging combined with Zipcode hybridization can distinguish cell populations. Purified CD4 and CD8 T mouse T cells were labeled with a caged anti-CD45 Ab-DNA conjugate and then spatially confined to separate tubes. ‘CD4 T cells’ were uniquely illuminated prior to addition of the first Zipcode (ZC1) to both tubes, followed by a washing step. ‘CD8 T cells’ were then illuminated and a second Zipcode (ZC2) was added to both tubes followed by washing. The contents of both tubes were pooled and introduced into the 10X scRNA-Seq pipeline. Nearest neighbor clustering of the populations sorted CD4 and CD8 T cell signatures from each other. We observed good (95%) correlation between dominant Zipcode identity and assignment to CD4 or CD8 clusters. We believe this error rate stems from the fact that the starting isolated populations were also approximately 95% pure by flow cytometry. **(a)** Flow cytometry plots of purified ‘CD4’ (population 1) and ‘CD8’ (population 2) populations used for the following study. CD4 and CD8 staining were used to define single positive cells with the % of cells in the single positive gate displayed. **(b)** Workflow for validation study. Population 1 was photo-uncaged w/ 365 nm light illumination prior to addition of Zipcode 1 while population 2 was illuminated prior to Zipcode 2 addition as a simulation of the workflow in tissue. **(c)** Histogram of (ZC1-ZC2)/(ZC1+ZC2) showing the bimodal distribution of cells with a ZC1 and ZC2 majority population as well as a small population centered about 0, suggesting presence of mixed doublets. Line at 0 denotes demarcation of cells into either ZC1 or ZC2 dominant population. (**d)** UMAP dimensional reduction of live cells with dominant Zipcode ID determined from (c) overlaid (blue: ZC1, red: ZC2) **(e)** Proportions of ZC1 and ZC2 dominant cells within each cluster. **(f)** Overlay of normalized gene expression levels for selected genes that define CD4 and CD8 T cells populations.

**Supplementary Figure 2:**
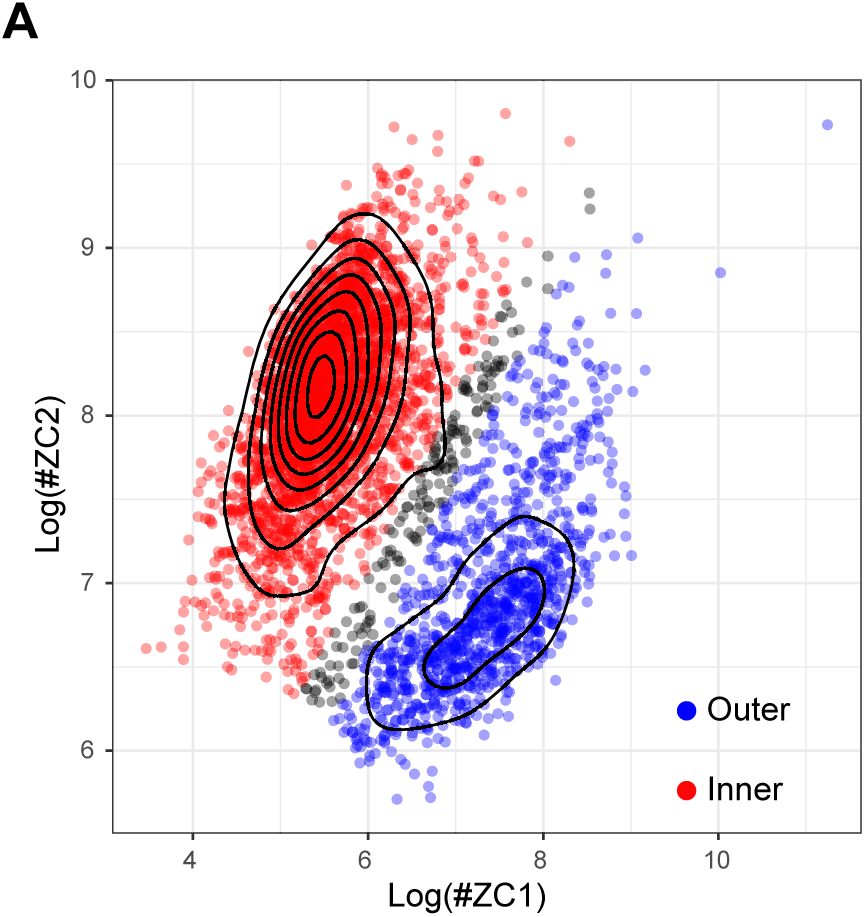
**(a)** Plot of Zipcode counts for Zipcode 1 and 2 on a logarithmic scale for cells detected using Cell Ranger defined cutoff. Regional calls were made based on dominance of ZC1 or ZC2 using k-means clustering to identify three groups. Cells within the cluster termed as Outer (Blue) had relatively more ZC1 counts and inversely, cells within the Inner (Red) cluster were dominated by ZC2 counts. Cells in-between were determined to be ‘ambiguous’ and excluded from downstream analysis.

**Supplementary Figure 3:**
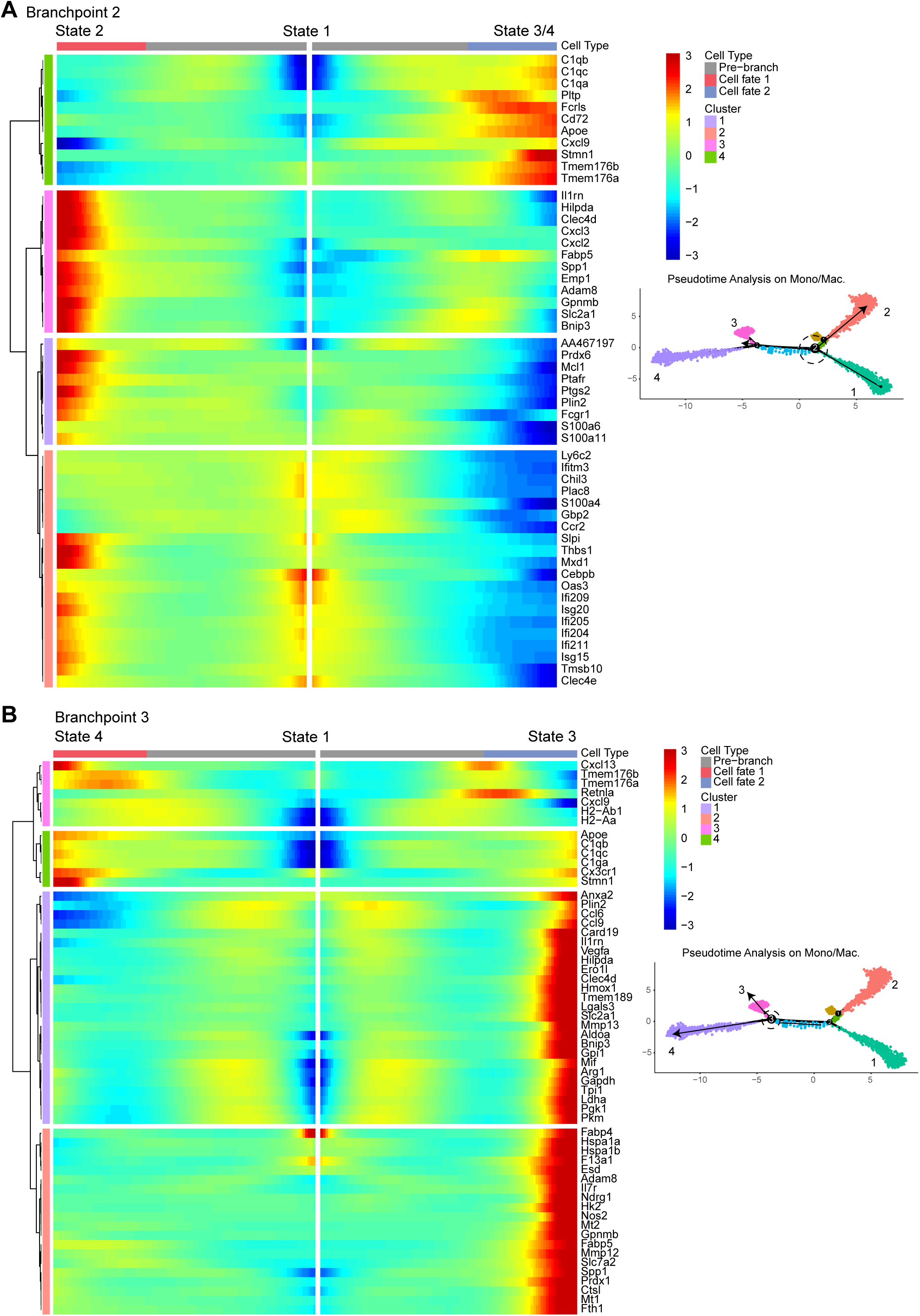
Heatmap of gene expression (color) over pseudotime (columns) for most significantly branch-dependent genes (rows) as determined by BEAM analysis. Genes are hierarchically clustered to illustrate genes that share expression patterns across branch point. **(a)** For the first branch point (from state 1 to either state 2 or 3/4) and **(b)** for second branch point differentiating states 3 and 4.

**Supplementary Figure 4:**
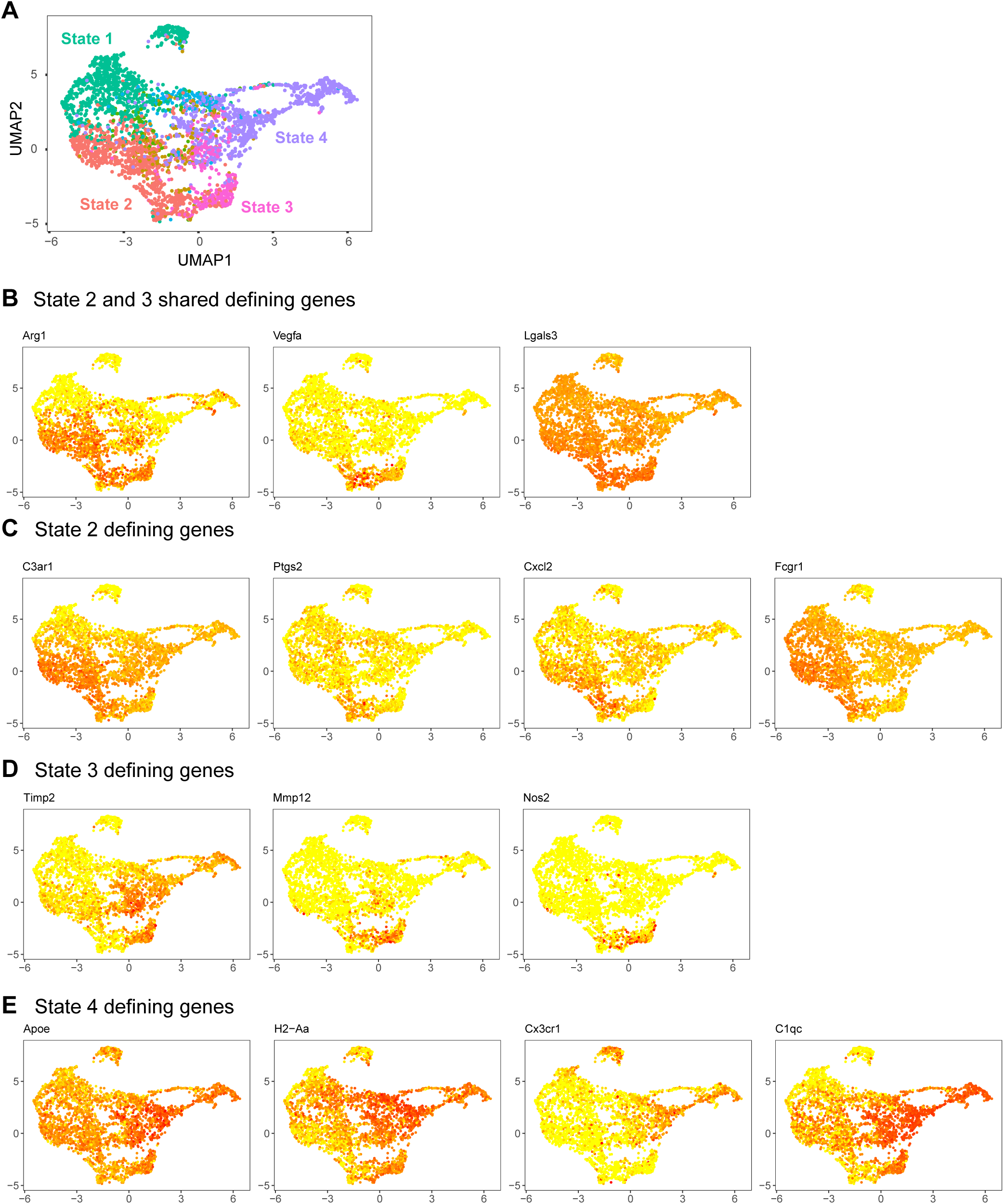
Feature plots of monocyte/macrophage population for selected genes that define terminal states 2-4. **(a)** UMAP dimensional reduction of monocyte/macrophage population with state identities overlaid. **(b)** Genes that are expressed by both states 2 and 3 over state 4. Despite the differentiation trajectory, States 2 and 3 shared overexpression of *Arg1* and angiogenesis factors (*Lgals3*, *Vegfa*) while under-expressing MHCII (*H2-Aa* and *H2-Ab1*) compared to State 4. **(c)** State 2 diverged from State 3 through expression of *C3ar1*, *Ptgs2, Cxcl2, Fcgr1* **(d)** Genes overexpressed in state 3. State 3 was defined by overexpression of genes such as *Nos2*, *Timp2*, and *Mmp12.* **(e)** Genes overexpressed in state 4. State 4 was defined by expression of *H2-Aa*, *C1qc/a*, *Apoe* and *Cx3cr1.* Color scale indicates log-normalized transcript counts.

**Supplementary Figure 5:**
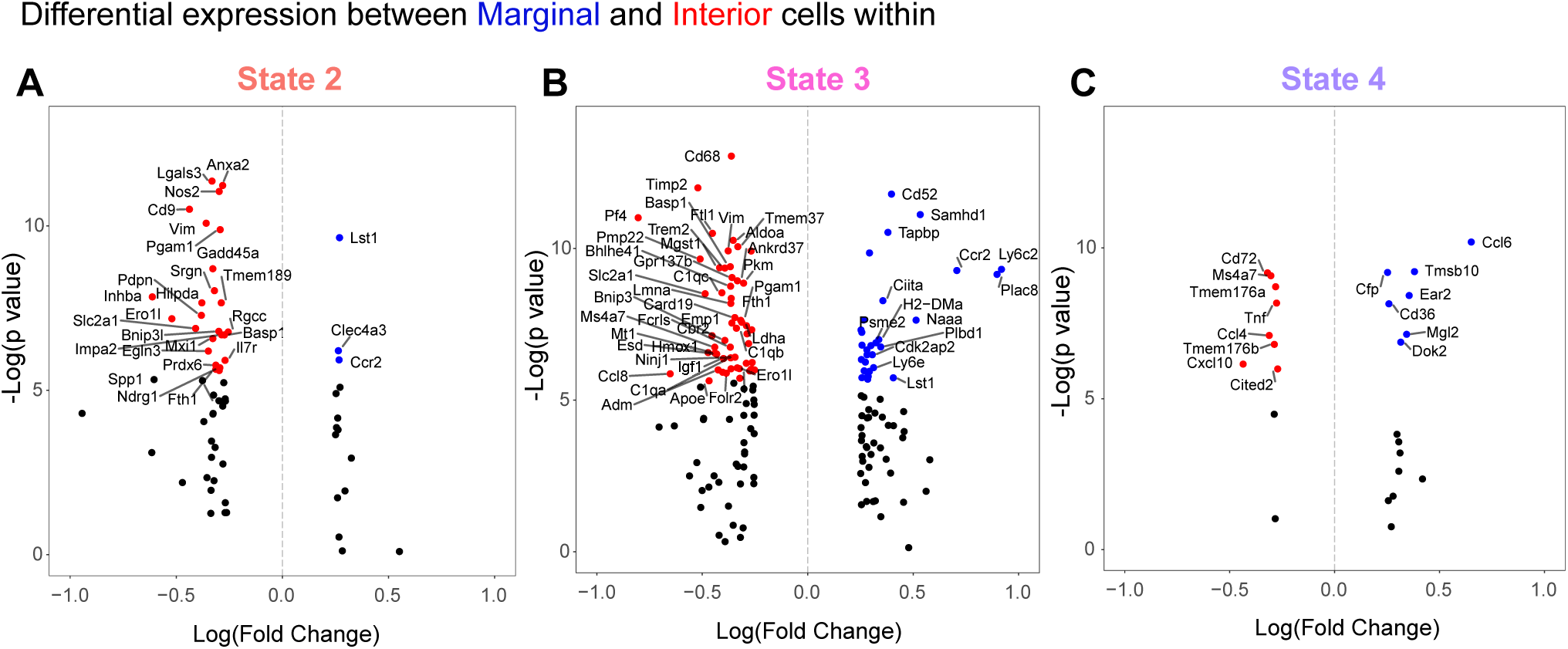
**(a-c)** Differential Gene expression analysis between margin and interior cells for each of the terminal macrophage states (2-4) defined through Monocle analysis as shown in Figure 4G. Volcano plots show genes with p-value (Bonferroni corrected) > 0.05 colored blue for margin enriched and red for interior enriched.

**Supplementary Figure 6:**
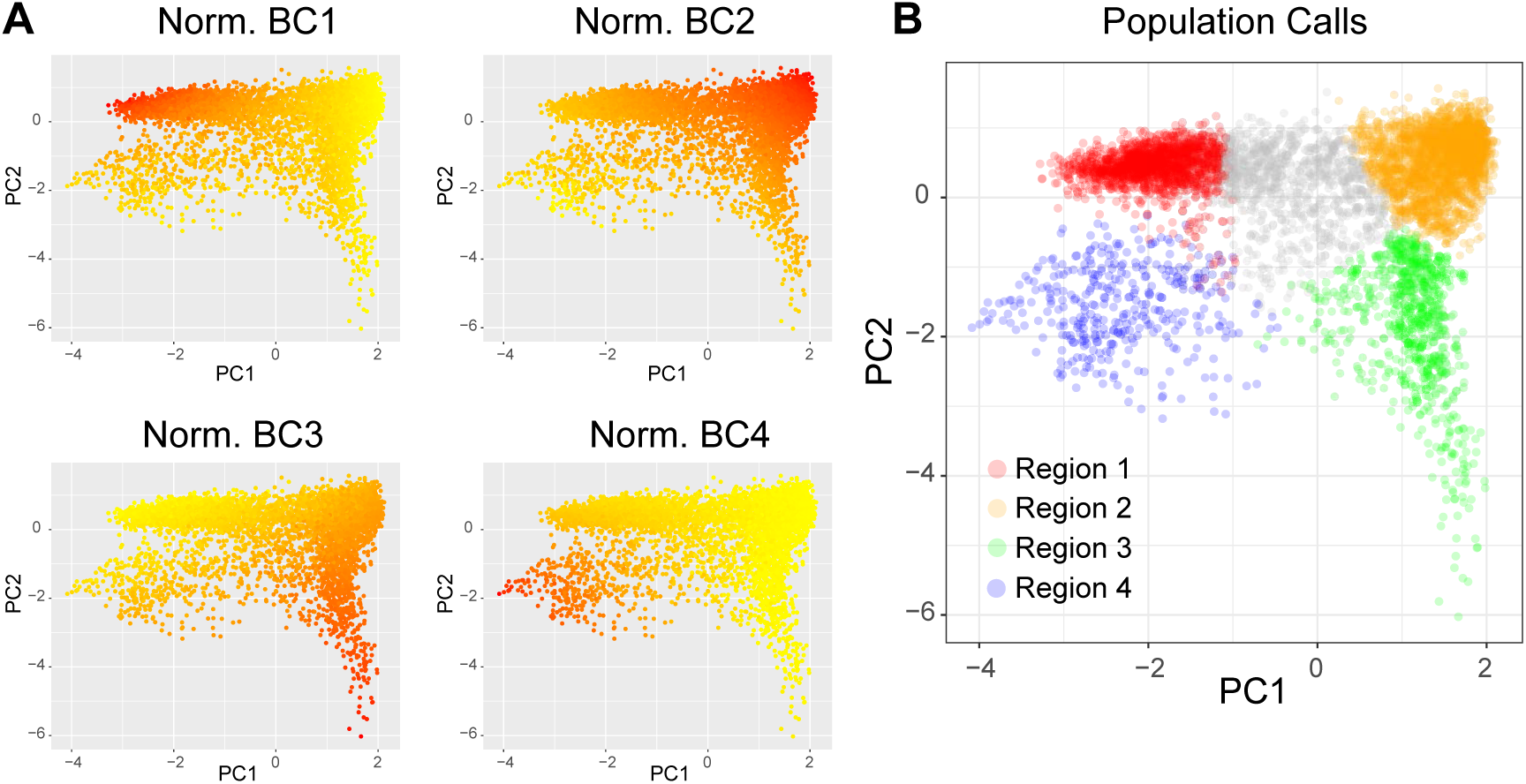
**(a)** Raw counts for Zipcodes 1-4 from live cells as defined by CellRanger were normalized to the total Zipcode count for each cell (i.e. % of ZC). Principal components were calculated using these 4 parameters. Plots show PC1 vs PC2 with the normalized ZC count for 1-4 overlaid. **(b)** k-means clustering was used to identify 5 clusters based on the normalized ZC counts in order to identify ZC1-4 dominant populations. Color overlays correspond to the regional call used in Figure 5.

**Supplementary Figure 7:**
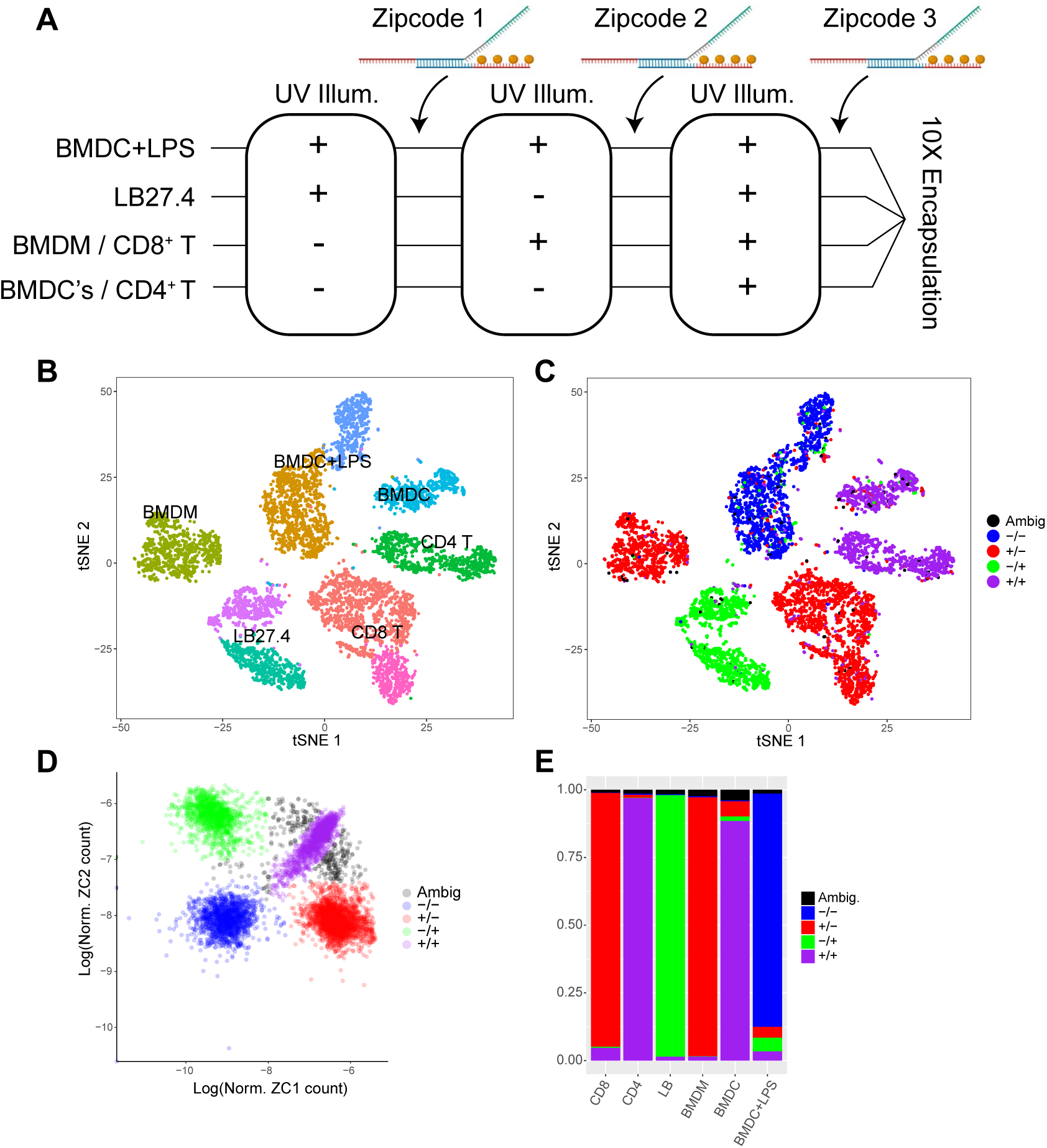
In-tube validation of combinatorial Zipcode labeling strategy **(a)** Isolated primary mouse immune cells of the designated types were labeled with the caged anti-CD45 anchor conjugate. Each population was then subjected to the specified series of illuminations and Zipcode additions with Zipcodes 1 and 2 used to determine identity and Zipcode 3 which was added onto all cells as a reference for normalization during analysis. Cell were then pooled and run through the modified 10X pipeline in order to recover both Zipcode and cDNA single cell libraries. **(b)** TSNE Plot showing major immune cell populations annotated. **(c)** Same tSNE plot with combinatorial Zipcode id overlaid. Black dots represent ambiguous Zipcode calls. **(d)** Plot of log-transformed, normalized Zipcode counts for ZC1 vs ZC2 (normalized to total ZC1-3 count) with Zipcode combination id’s overlaid. **(e)** Bar charts showing proportion of cells with a particular Zipcode combination for a given cell type. Colors consistent with those used in (b).

**Supplementary Figure 8:**
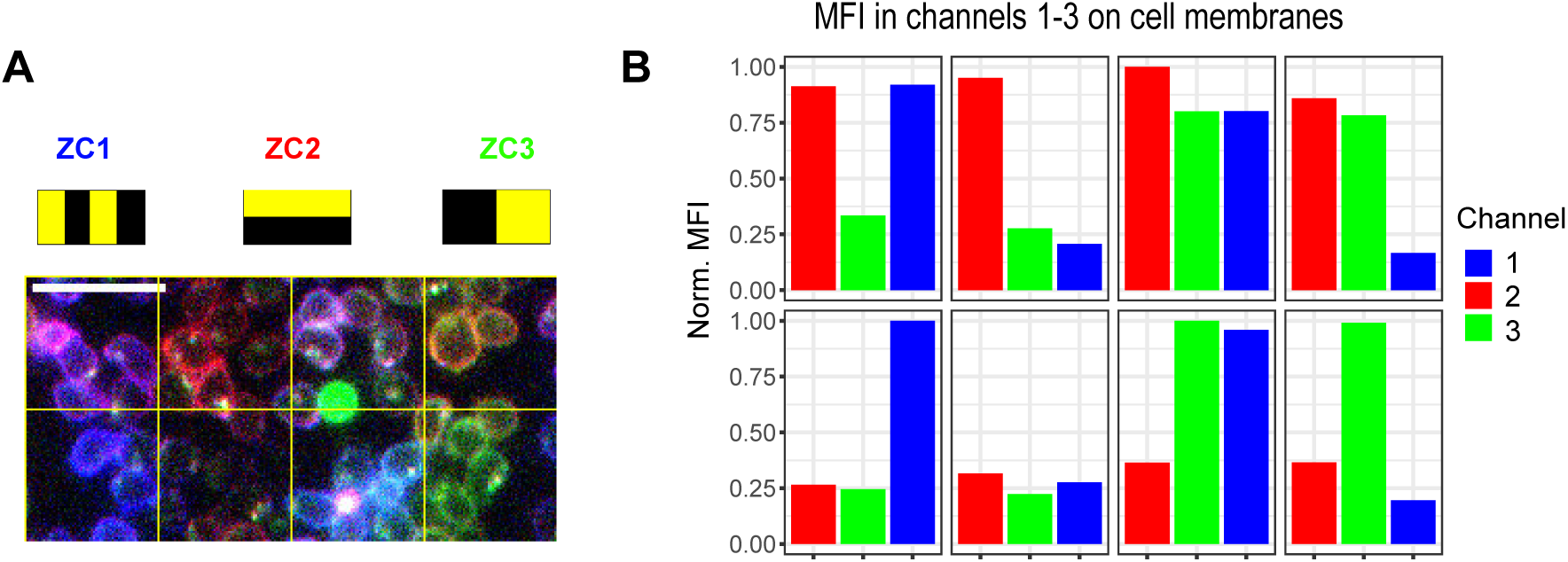
8 region Zipcode delineation using combinatorial design with 20-micron resolution. **(a)** Microscope image of field of mouse T cells following 3 rounds of illumination w/ patterns shown and addition of three distinct Zipcode species with conjugated fluorophores. Experimental setup was similar to Figure 5I. Each grid square is approximately 20 × 20 µm. **(b)** Quantification of mean fluorescence intensity for each channel (Zipcode) on cell surfaces. Briefly, for each grid square, cell membranes were defined using the union of masks generated by thresholding in each channel. Mean channel intensity was calculated for pixels within this combined mask and background intensity for each channel was subtracted. Bar charts are shown normalized to maximal MFI in all 8 grid squares. Scale bar = 20 µm.

